# Mitochondrial genome rearrangements and virulence-associated features in a clinical isolate of the emerging pathogen *Wickerhamomyces anomalus*

**DOI:** 10.1101/2025.08.29.673123

**Authors:** M Cáceres-Valdiviezo, L Salvatierra- Chung Sang, JC Tapia, Y Acosta, Andres Caceres-Valdiviezo, G Morey-León, JC Fernández-Cadena, J.A. Ugalde, D Andrade-Molina

## Abstract

**Background:** *Wickerhamomyces anomalus,* also known as *Candida pelliculosa,* traditionally regarded as an environmental yeast, has emerged as an opportunistic yeast implicated in invasive infections and hospital outbreaks, especially among neonatal and immunocompromised individuals. Despite its clinical relevance, the genomic determinants underlying the pathogenesis and virulence remain poorly understood.

**Results:** Here, we report the first high-quality nuclear and complete mitochondrial genome assemblies from a clinical isolate of *W. anomalus.* The nuclear genome spans 13.9 Mb with 34.5% GC content and contains 6,305 protein-coding genes, 8,235 coding sequences, and 147 tRNAs, with 98.1% completeness by BUSCO. The mitochondrial genome (47.6 kb) encodes 19 protein-coding genes, 26 tRNAs, and two rRNAs. Remarkably, it exhibits atypical rearrangements, including inverted repeats and atp9 duplications.

**Conclusions:** We provide the first de novo nuclear genome from a clinical *W. anomalus* isolate, along with the first complete mitochondrial genome for this species, establishing a high-quality reference for this emerging pathogen. The genomic innovations, together with a high content of repetitive and non-coding elements, suggest enhanced plasticity that may underpin metabolic flexibility and survival under host-associated stresses. These rearrangements represent an underappreciated mechanism of adaptation in emerging fungal pathogens, contributing to the clinical relevance of *W. anomalus.* These genomic resources facilitate rigorous comparative analyses to indentify determinants of pathogenicity, virulence, and antifungal resistance, accelerating the development of improved diagnostics, surveillance, and management strategies for *W. anomalus* infections

## Background

Fungal infections are responsible for more than 1.5 million deaths per year, with candidiasis the most common fungal disease in hospitals (1–3). Although *Candida albicans* is the most common cause of invasive candidiasis, non-*albicans* candidiasis cases are increasingly reported worldwide, surpassing *C. albicans* candidiasis in some cases (1,4). Among the non-albicans *Candida* species, *Wickerhamomyces anomalus* (previously classified as *Pichia anomala*, *Hansenula anomala,* and *Candida pelliculosa*) has progressively emerged as a concerning pathogen in nosocomial outbreaks among neonates and immunocompromised patients (5).

*W. anomalus* is a diploid yeast isolated from diverse habitats, including plants, insects, soil, human tissues, and marine environments. Its wide metabolic and physiological diversity enables it to thrive over diverse environmental conditions, such as pH and extreme temperature fluctuations, and osmotic stress (6–8). These characteristics render *W. anomalus* a versatile organism, widely employed in the food, environmental, industrial, and medical sectors (9–11). Metabolism and metabolic adaptation provide essential biosynthetic precursors and energy for critical biological functions in yeast (12). Mitochondria have a crucial role in coordinating diverse anabolic and catabolic pathways fundamental to energy generation, metabolism, and pathogenesis in yeast (13). In recent years, there has been a concentrated interest in the structural and functional characteristics of mitochondria in pathogenic yeasts such as *C. albicans* (14) and *C. auris* (15), while the mitochondrial genome of *W. anomalous* is unknown. Research on the evolution and functionality of the mitochondrial genome can be challenging in certain fungal species, primarily due to notable difficulties associated with *de novo* assembly and intron prediction within the protein-coding genes (PCGs) (16–18). Moreover, increasing evidence suggests that mitochondrial genetic features may contribute to the ability of specific yeasts to survive in hostile host environments, evade immune responses, and establish ongoing infections (19).

This knowledge gap is relevant in the case of *W. anomalus,* an emerging opportunistic pathogen associated with several clonal outbreaks worldwide. These infections are linked to poor clinical outcomes, with reported mortality rates exceeding 40% in some cases (20–22). Moreover, clinical isolates of *W. anomalus* are increasingly exhibiting resistance to azoles, including fluconazole (23,24). Despite its clinical relevance, the relationship between the genomic structure and its virulence and pathogenicity has not been previously studied.

Intending to improve our knowledge on this opportunistic yeast, we present the first nuclear and mitochondrial genome assembly of a clinical isolate of *W. anomalus* using short and long reads generated by Illumina and PacBio sequencing, respectively. Our findings indicate that the genome of *W. anomalus* is highly congruent, with high levels of similarity to available non-clinical genomes. Analysis using genomes of *Candida* spp. and *Saccharomyces cerevisiae* as reference revealed a set of orthologous proteins such as synthetases, oligopeptide transporters, cell wall-associated genes, and antioxidant enzymes, which may play a role in the virulence of *W. anomalus*. However, the majority of the genome remains uncharacterized.

Further studies are required to decipher the role of these hypothetical proteins in *W. anomalus*, as they might contribute to species-specific traits that could play a role in its pathogenesis.

## Results

### Isolate resistance profile

The *in vitro* antifungal susceptibility test showed sensitivity to fluconazole, voriconazole, caspofungin, micafungin, amphotericin B, and flucytosine due to the low off–scale MIC value of Vitek 2 AST YS08 (Supplementary table 1).

### Assembling the nuclear *W. anomalus* genome

A combination of long-read PacBio and short-read Illumina sequencing was employed to obtain the genome of the *W. anomalus* strain isolated from a clinical sample (hereafter CpEC_Uees). Before assembly, the long PacBio subreads were pre-processed to generate Highly Accurate Single-Molecule Consensus reads (ccs). Different genome assembly algorithms were employed to create high-quality and high-coverage assemblies. The sequencing reads were assembled using the hybrid assemblers Wengan (25) and MaSuRCA (26), as well as the long-read assemblers Flye (27) and SMARTdenovo (28). The resulting assemblies were polished thrice with the short Illumina reads to remove sequence and structural errors. Traditional assembly metrics such as contig count, maximum contig size, N50, and genome size were used to compare the resulting assemblies (Table 1). Among these, the CpEC_Uees genome generated by MaSuRCA had the highest contiguity, with 20 contigs of maximum length up to 2.95 Mbp, N50 of 1.28 Mbp, and a GC content of 34.51%.

**Table 1.** Summary information of the assemblies generated with the different assemblers used.

|  | **Wengan** | **MaSuRCA** | **Flye** | **SMARTdenovo** | **Canu-SMARTdenovo** |
| --- | --- | --- | --- | --- | --- |
| **Contig Numbers** | 342 | 20 | 52 | 30 | 29 |
| **Total genome size (pb)** | 15,641,129 | 13,991,713 | 14,001,624 | 14,097,927 | 14,004,920 |
| **GC content (%)** | 34.63 | 34.51 | 34.69 | 34.53 | 34.56 |
| **N50 (bp)** | 74,133 | 1,283,267 | 1,422,051 | 891,120 | 960,118 |
| **N75 (bp)** | 39,304 | 1,112,314 | 566,893 | 583,755 | 576,198 |
| **Largest Contig** | 294,006 | 2,946,481 | 2,746,987 | 3,632,912 | 3,626,365 |
| **BUSCO score completeness** | 67% | 98.1% | 96.4% | 88.7% | 96.6% |

The MaSuRCA contig set was then scaffolded using RagTag (29) and the previously published chromosome-level genome (GCA_019321675.1) as reference. Three contigs (scf7180000000186, scf7180000000191, scf7180000000194) comprising ∼90kb did not align to the reference genome and were further processed (Supplementary Figure 1). The final CpEC_Uees draft assembly yielded 10 scaffolds, with an N50 of 2,125,767 bp and a total length of 13,959,047 bp. Moreover, out of the 10 scaffolds, seven correspond to Chromosomes 1-7, and one to the mitogenome.

The completeness of the genome assembly was assessed using the BUSCO tool (30), which evaluates the presence of universally conserved single-copy genes. BUSCO identified 2,098 out of 2,137 genes (98.1%) of the lineage-specific saccharomycetes group in the CpEC_Uees assembly as complete, with an additional fragmented gene. These results are comparable to those of the available assemblies of *W. anomalus* at NCBI. Additionally, more than 98% of PacBio and Illumina data were mapped to the final assembly with coverage of an average depth of 190X and 95X, respectively (Supplementary Table 2).

To evaluate the structural rearrangement of the CpEC_UEES genome, we aligned the draft assemblies from MaSuRCa and Canu-SMARTdenovo against the GCA_019321675.1 reference genome (Supplementary Figure 1). A significant macrosynteny and high sequence homology (>95% sequence identity) were observed. Although the MaSuRCa assembly contained 10 strand matches (purple lines) and 4 strand inversions (light blue lines) (Figure 1B), only two inversions were also present in the Canu-SMARTdenovo assembly (Supplementary Figure 1). Overall, these findings reveal that the structural rearrangement of the CpEC_UEES genome is largely conserved in comparison with the reference genome.

**Figure 1.**
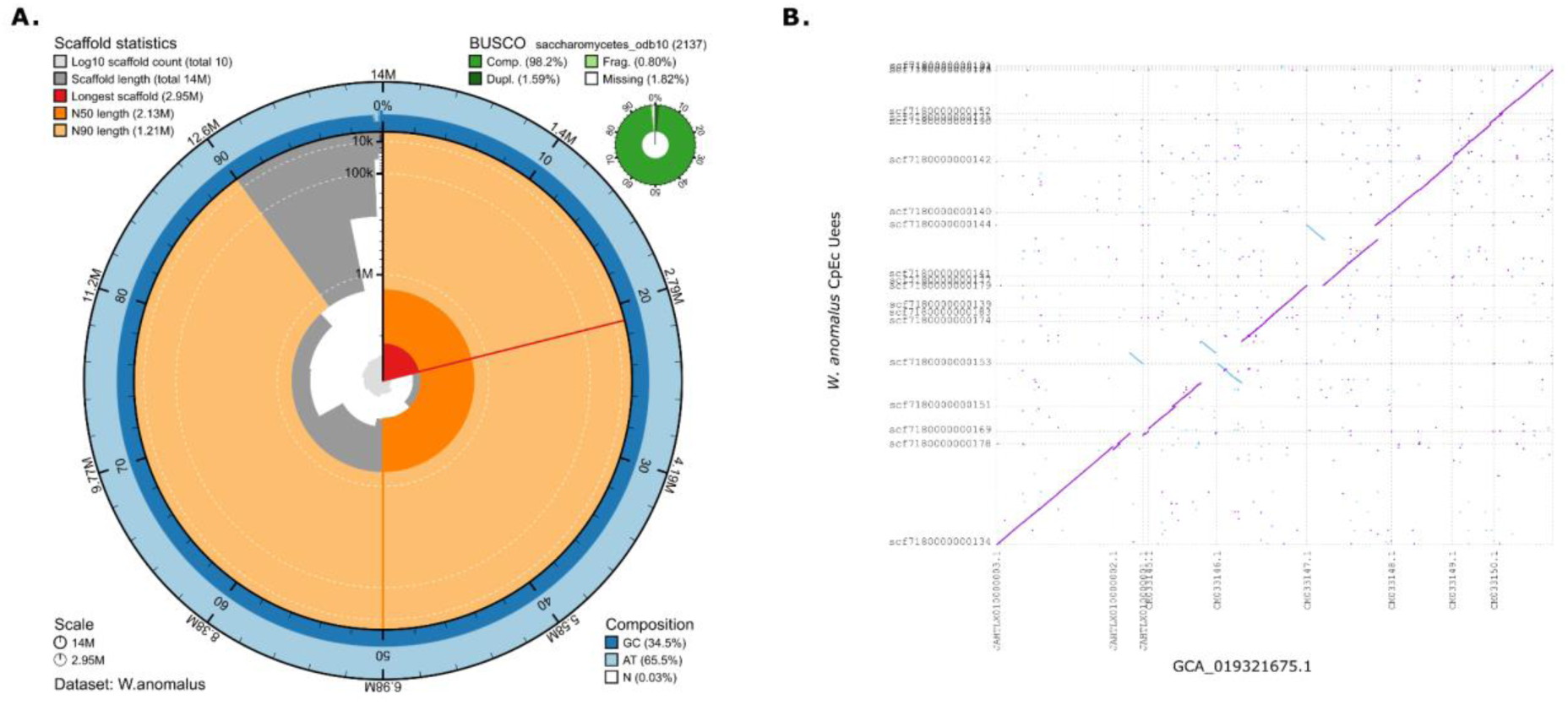
Genome assembly statistics and a dot-plot depicting the scaffold-level and contig-level assemblies, respectively, compared with the reference genome GCA_019321675.1. **A.** Circular plot visualising the assembly metrics of the final *W. anomalus* draft genome assembly using BlobToolKit viewer (31). **B.** The Dot-plot was generated using MUMmer 4.0.0 (NUCmer) (32). Diagonal lines represent alignment between the reference genome (GCA_019321675.1) (horizontal) and CpEC_UEES (contigs, vertical). Purple lines indicate high similarity between strands, while light blue lines represent inverse alignment. Points near the diagonal indicate the co-linearity of genome sequences.

### CpEC_UEES genome annotation

Protein-coding genes were annotated using a *de novo* homology-based approach. A total of 6,324 putative genes were predicted, of which 5,305 genes (84%) were functionally annotated by eggNOG-mapper (33). The draft assembly was also used to identify 173 tRNAs and 11 truncated transposable elements belonging to the Ty1, Ty3, and Ty5 transposon families.

From the 6,324 predicted proteins, 4,451 sequences (∼70.4%) were annotated with GO terms and placed in three main domains: Molecular Function, Cellular Component, and Biological Process (Figure 2a, Supplementary Table 3). Notably, a substantial number of genes were related to cellular anatomical entities (4,266 sequences) and Cellular processes (4,083 sequences) within the Cellular component and Biological process domains, respectively. The Molecular function domain revealed a predominant presence of genes associated with catalytic and ligand-binding activity.

**Figure 2.**
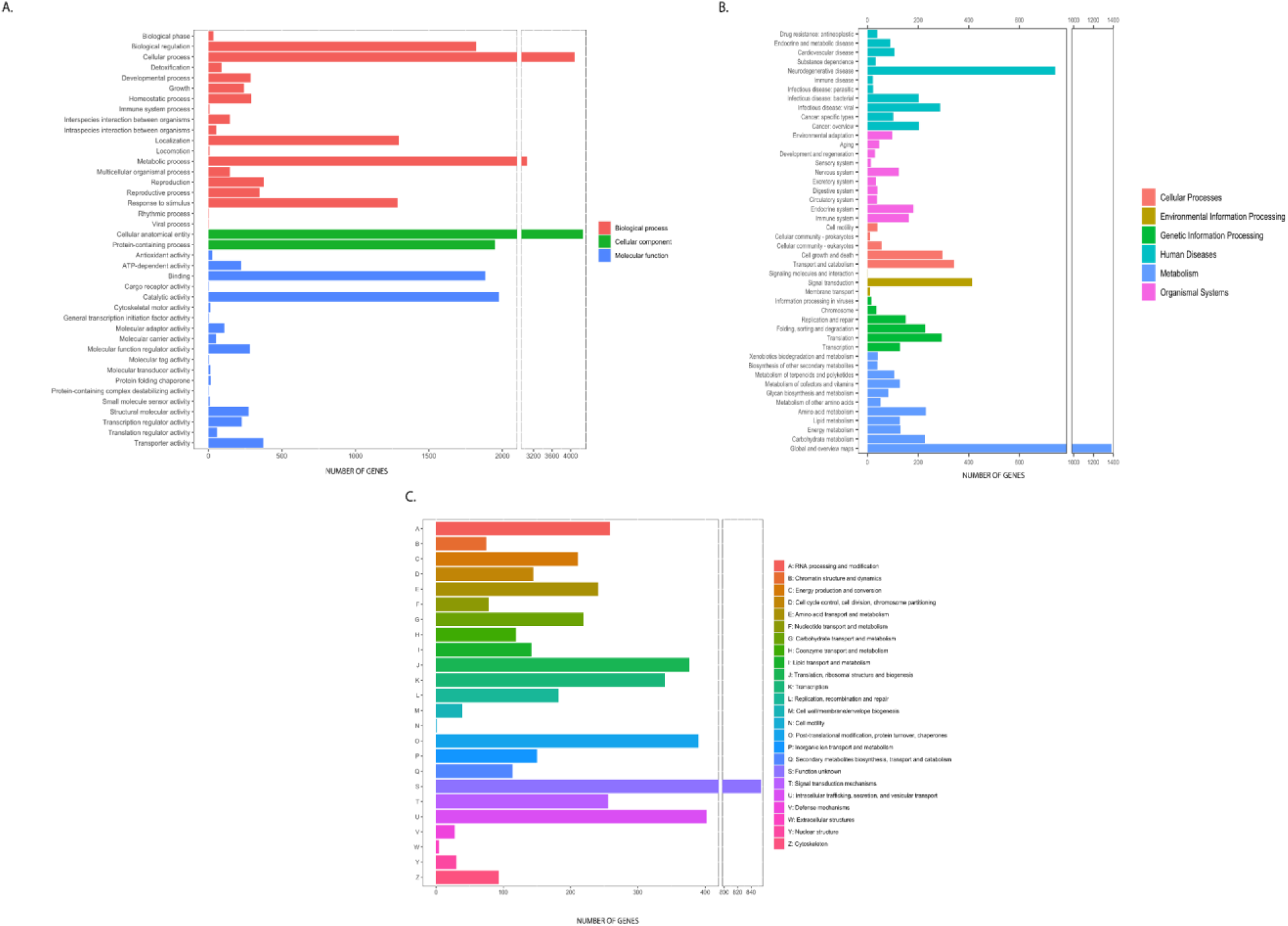
Distribution of Gene Ontology (GO), Eukaryotic Orthologous Groups (KOGs), and Kyoto Encyclopaedia of Genes and Genomes (KEGG) annotations of predicted proteins from *W. anomalus* CpEC genome. **A.** Functional distribution of the predicted proteins according to three major hierarchical GO terms: Biological process (red), Cellular component (green), Molecular function (blue). **B.** Distribution of KOG annotation profiles. **C.** Classification and functional distribution of the predicted proteins according to six major hierarchical KEGG terms.

In parallel, 5,197 genes were classified into KOG (euKaryotic Orthologous Groups) families across 24 categories. The most abundant classes were U (Intracellular trafficking and secretion), J (Translation ribosomal, structure, and biogenesis), O (Post-translational modifications, protein turnover, chaperones), and K (Transcription) (Figure 2b, Supplementary Table 4). Furthermore, KEGG annotation classified 3,156 protein-coding sequences (45.6%) into 396 pathways (Figure 2c, Supplementary Table 5). Among these, the metabolic pathways (KO:01100) were the most enriched KEGG term with 638 genes, followed by biosynthesis of secondary metabolites (KO:01110, 261 genes), amyotrophic lateral sclerosis (KO:05014, 132 genes), and pathways of neurodegeneration - multiple diseases (KO:05022, 126 genes).

### Mitogenome assembly

Of the three unaligned contigs, we identified a single contig (scf7180000000186) corresponding to the mitochondrial genome based on strong homology to mitochondrial genes of *S. cerevisiae* (RefSeq: GCF_000146045.2)*, N. glabratus* (RefSeq: GCF_010111755.1), and *K. marxianus* (RefSeq: GCF_001417885.1). The final mitogenome assembly of *W. anomalus* consists of a single circular contig of 47,621 bp (GenBank ID: JBHVOA000000000) and a low GC content of 20.8%. It encodes 19 protein-encoding genes (PCGs), 26 tRNAs, and 2 rRNAs (Figure 3). Among these 47 genes, 27 are encoded by the leading strand, while the remaining 20 genes, including 9 protein-coding genes (*atp6, atp8, atp9, cox1, cox3, nad1, nad6, rps3,* and *lagli*), and 11 tRNAs (*trnA, trnC, trnD, trnE, trnG, trnI, trnN, trnR, trnS1, trnS2,* and *trnY*) are encoded by the lagging strand (Figure 3). The genome contains three group I introns (two of which belong to the subtypes IA3 and ID) totaling 4,634 bp (9.73% of the genome), of which 3,039 bp (65.58%) correspond to LADLIDADG (LD) ORFs (open reading frame). Two large inverted repeats (∼7kb each) with 100% identity were also identified, each harboring a single copy of *atp9* and *trnR* genes (Figure 3B). These repeats are located between positions 12,815 - 20,058 bp and 30,660 - 37,903 bp and are supported by long PacBio reads spanning the entire region, discarding assembly artifacts (Supplementary Figure 2).

**Figure 3.**
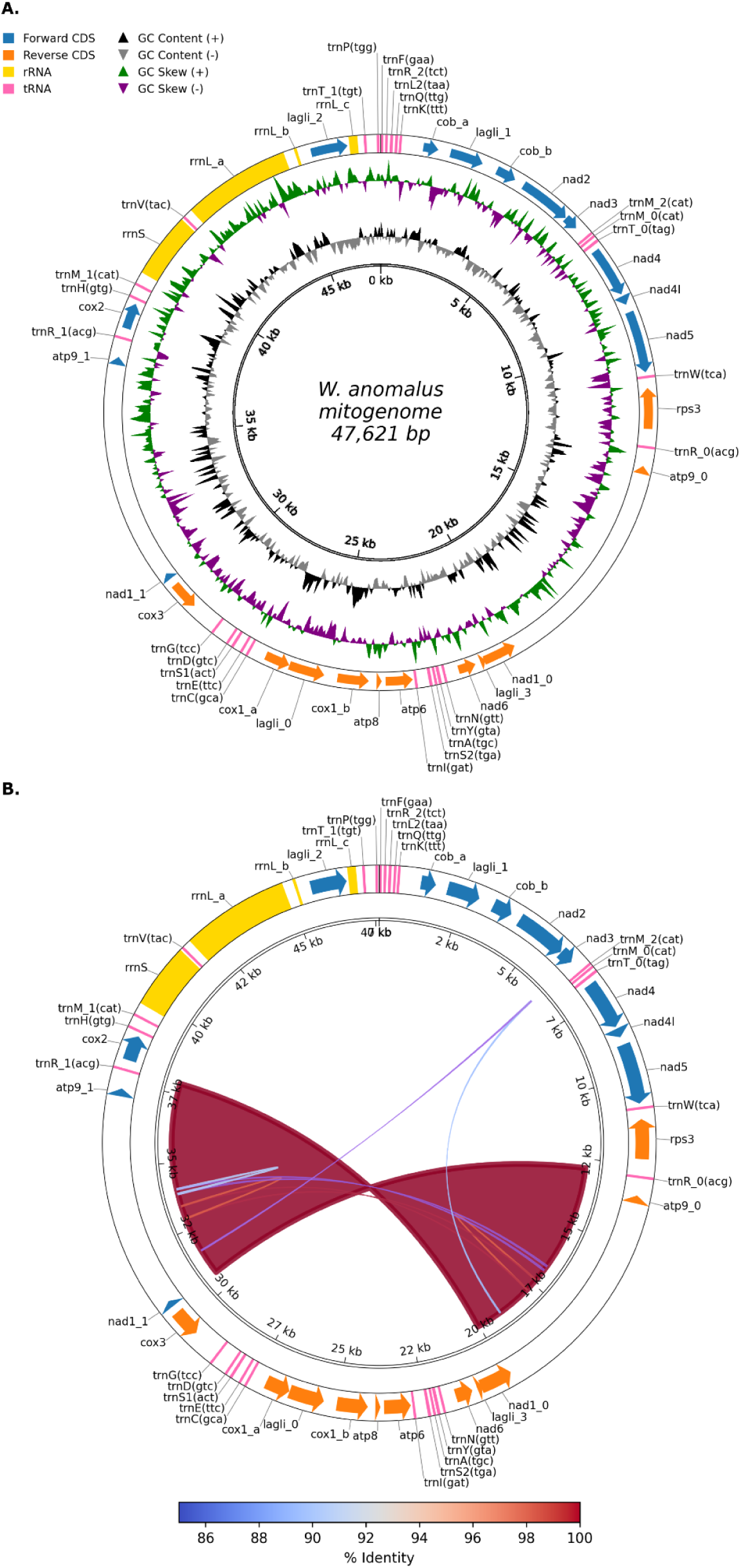
Circular representation of *W. anomalus* mitochondrial genome. **A.** The outer track depicts annotated genomic features, including 16 core mitochondrial protein-coding genes (*atp6, atp8, atp9, nad1-nad6, nad4l, cob, cox1-cox3*, and *rps3*), 26 tRNA genes, the ribosomal RNA subunits (*rrnS* and *rrnL*), and three LAGLIDADG (LD)-containing endonuclease genes. Track 2 and Track 3 show GC content and GC skew, respectively. **B.** Self-alignment of the mitochondrial genome, showing regions with e-value < 1e-10 and sequence identity > 85%. Two ∼7kb regions exhibit 100% identity, corresponding to inverted repeats that include *atp9* and *trnR*.

### Protein-coding genes in the mitogenome

The total length of the 19 PCGs is 19,755 bp, accounting for 41.48% of the entire mitochondrial sequence of *W. anomalus.* Of these, 16 correspond to the core mitochondrial genes (*atp6, atp8, atp9, cob, cox1, cox2, cox3, nad1, nad2, nad3, nad4, nad4l, nad5, nad6,* and *rps3*), with two copies of *atp9* being present.

The remaining three genes code for proteins exhibiting similarities to the LD homing endonucleases (HEs). Two of these were identified by both MITOS2 (34) and MFannot (35) and are located within the *rrnL* and *cox1* genes. The third ORF, located within an intron of the *cob* gene, was predicted only by MITOS2 (Table 2). However, this intronic ORF is considerably long (>900bp) and contains a well-conserved LD motif, suggesting that it is a true intron-encoded endonuclease. MITOS2 also predicted an additional short ORF (∼140 bp) containing part of the LD motif. However, BLAST analysis showed strong similarity (E-values < 4e-93) to *nad1* sequences from multiple fungal species. Consistent with this, MFannot re-annotation placed this sequence within the *nad1* coding region, and no Pfam domains associated with LD motifs were detected. We therefore annotated this region as part of the *nad1* gene. Most PCGs start with the canonical translation initiation codon ATG, except for the LD-encoding genes, which are initiated with ATA. Two genes (*cox2* and *rrnL*) use TAG as the stop codon, while the remaining genes use TAA. Codon usage analysis indicated that the most frequently used codons were TTA (Leu2), ATA (Met), AAT (Asn), TAT (Tyr), ATT (Ile), and TTT (Phe) with 718, 570, 471, 401, 311, and 230 counts, respectively. In contrast, the least commonly used codons were CGG (Arg), AGG (Arg), and CGC (Arg), each occurring once. Moreover, Relative Synonymous Codon Usage (RSCU) values revealed that ACT (4.05, Thr), AGA (3.44, Arg), and ACA (2.54, Thr) were used at disproportionately high frequencies relative to synonymous alternatives (Supplementary Table 6).

**Table 2.**
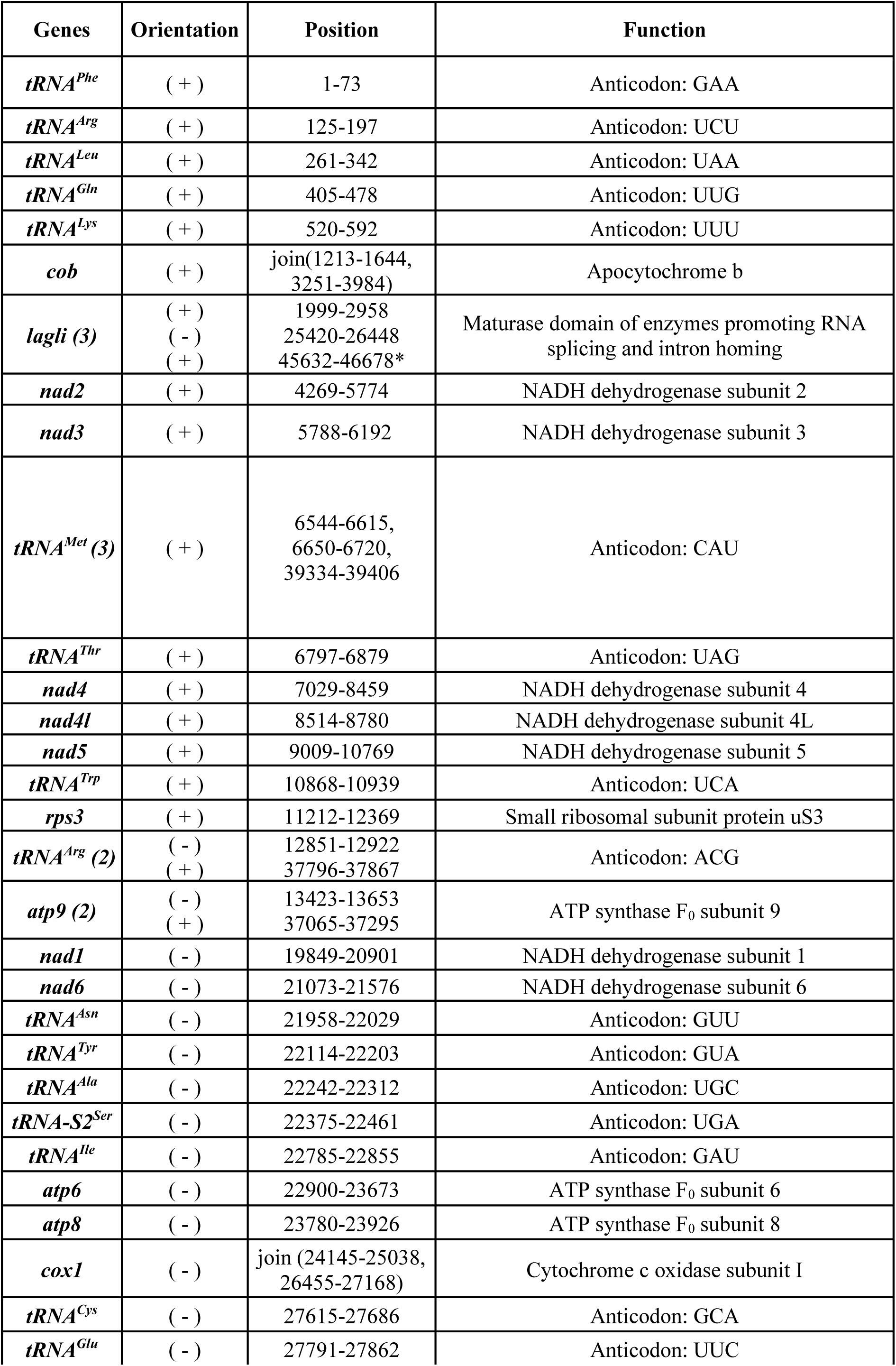
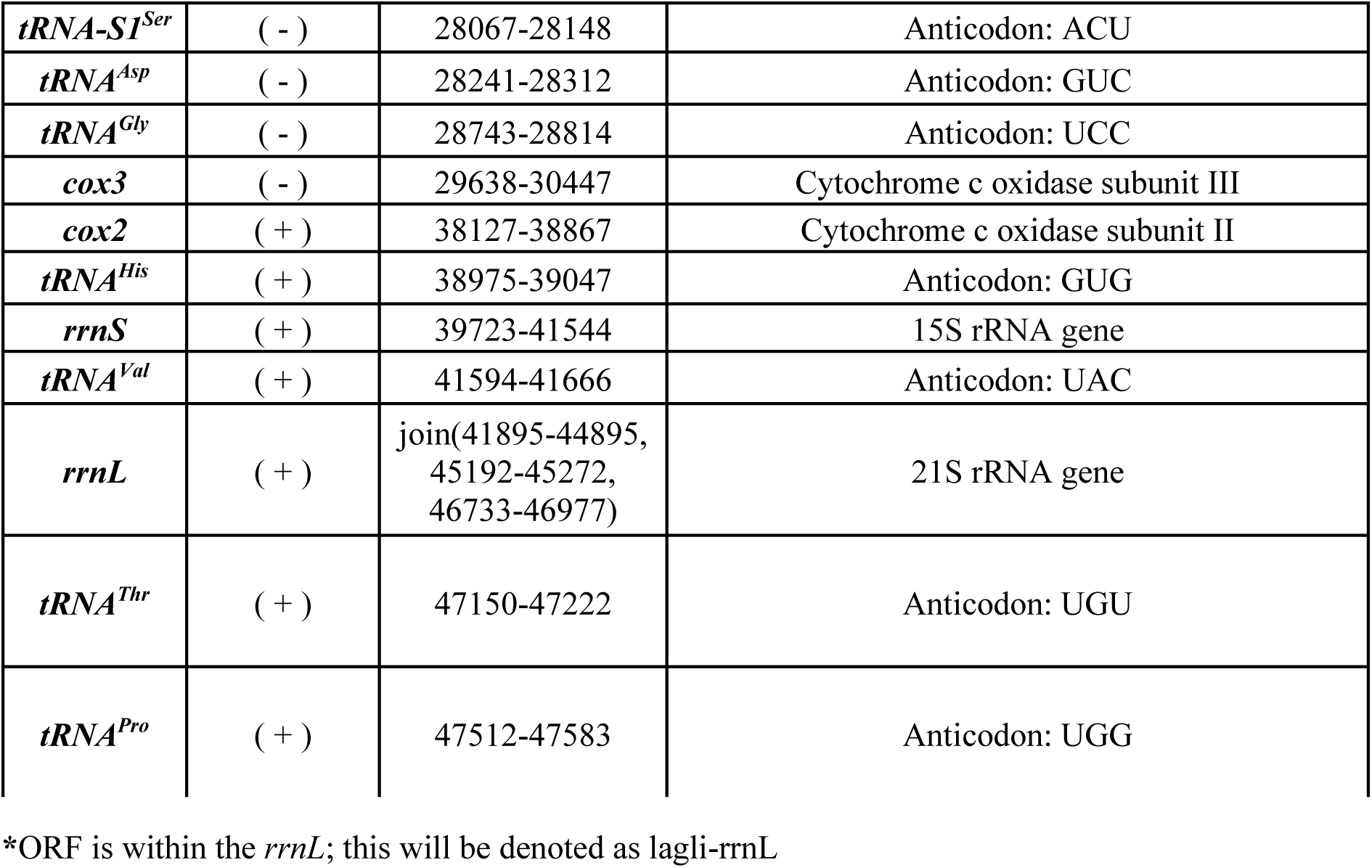
Annotation of the *W. anomalus* mitochondrial genome.

**Table 3.**
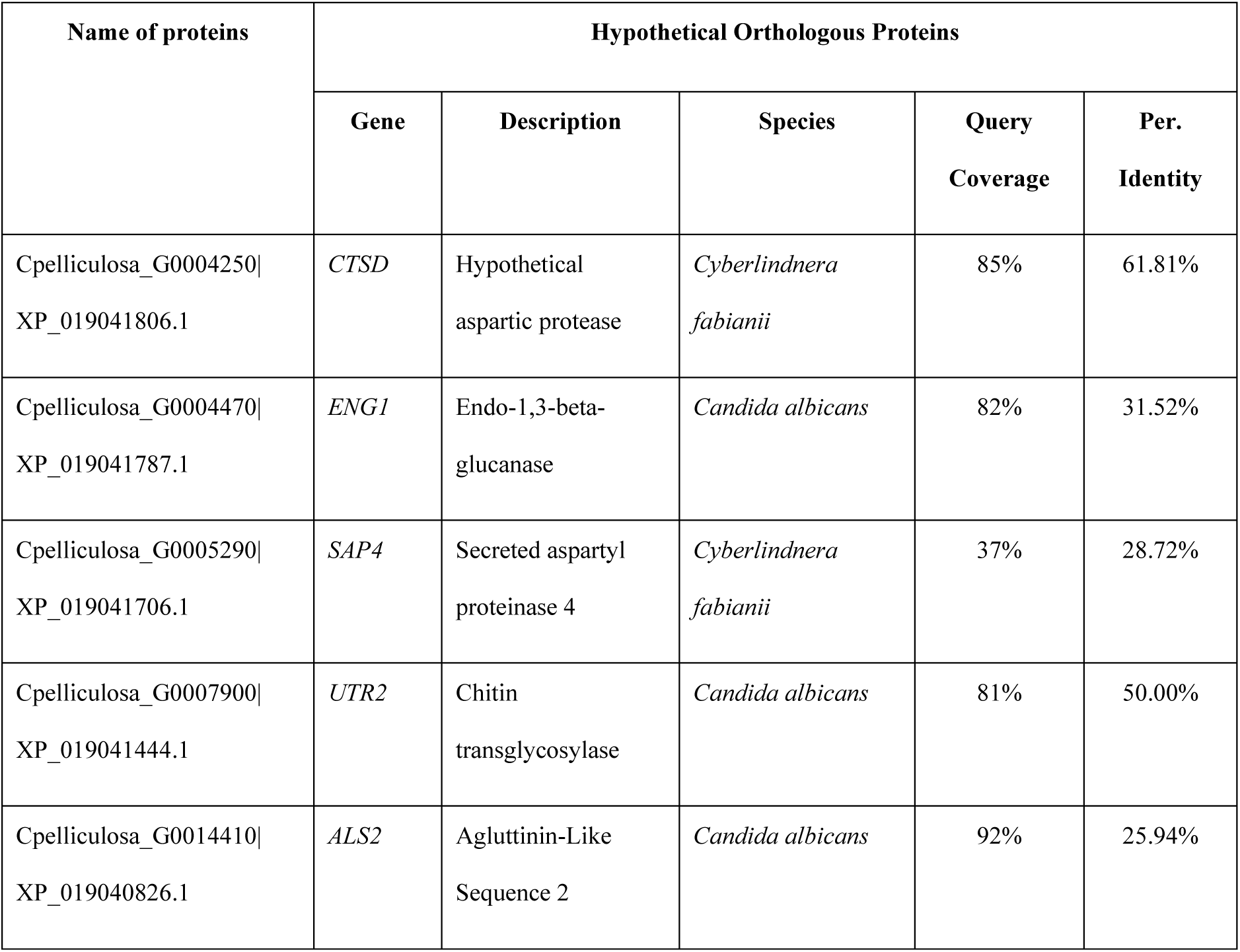
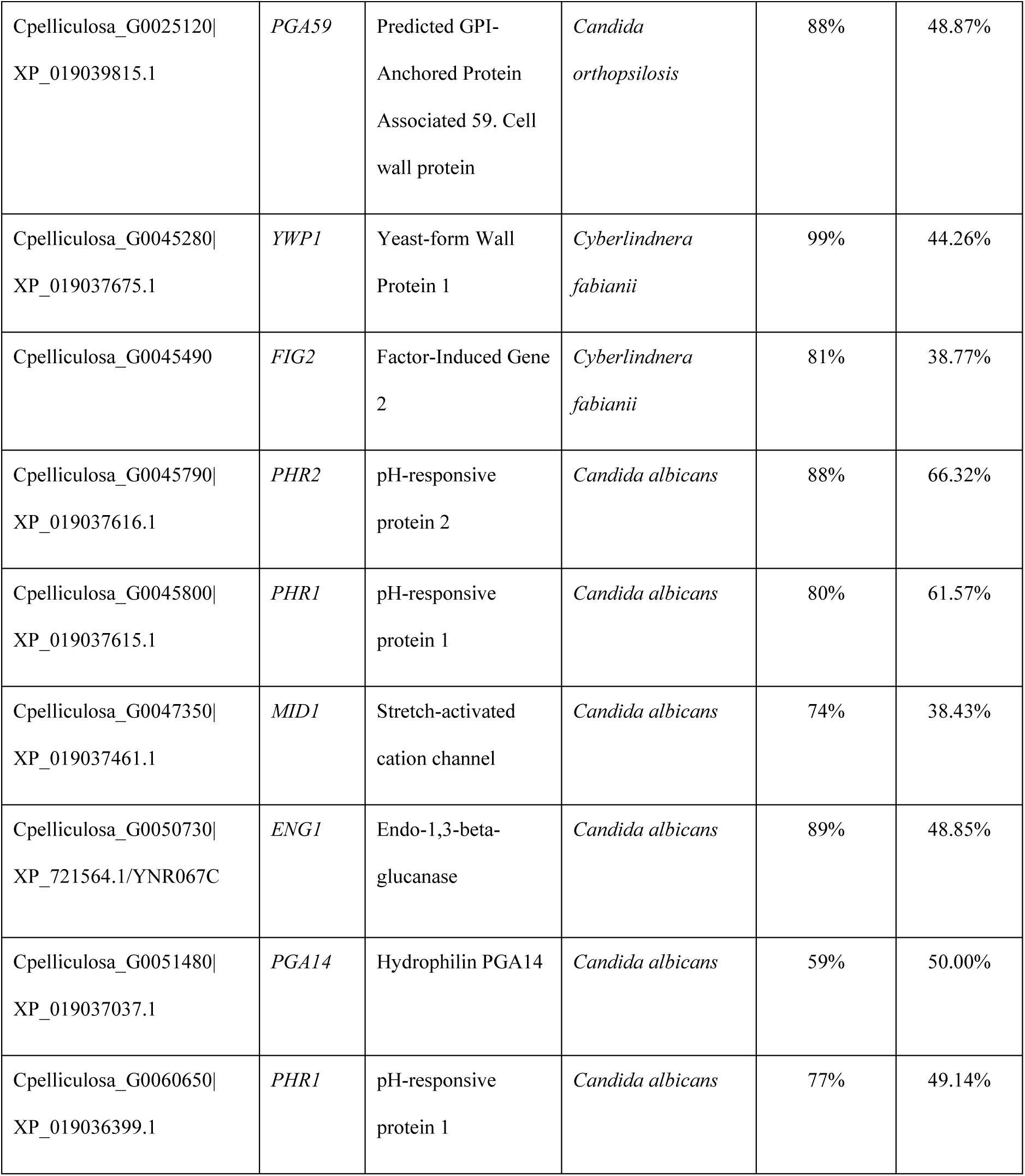
Hypothetical orthologous proteins of the predicted adhesin-like protein sequences.

### tRNA and rRNA genes

The *W. anomalus* mitochondrial sequence contains a total of 26 tRNA genes with a total length of 1,942 bp, accounting for 4.08% of the entire mitogenome. All of the tRNAs exhibit the characteristic clover-leaf structure, and their average length ranged from 71 bp (*trnM, trnA, and trnI*) to 90 bp (*trnY*), which was mainly due to the variable arm (Figure 4). There is one tRNA for each amino acid with additional isoforms for the amino acids Methionine (Met), Arginine (Arg), Serine (Ser), and Threonine (Thr) (Table 2). The mitogenome harbours three copies of the *trnM* (CAT) and *trnR* genes (two copies recognizing ACG, and one copy recognizing TCT), and two copies of the *trnS* (*trnS1* [ACT] and *trnS2* [TGA]) and the *trnT* genes (*trnT* [TAG], and *trnT* [TGT]).

**Figure 4.**
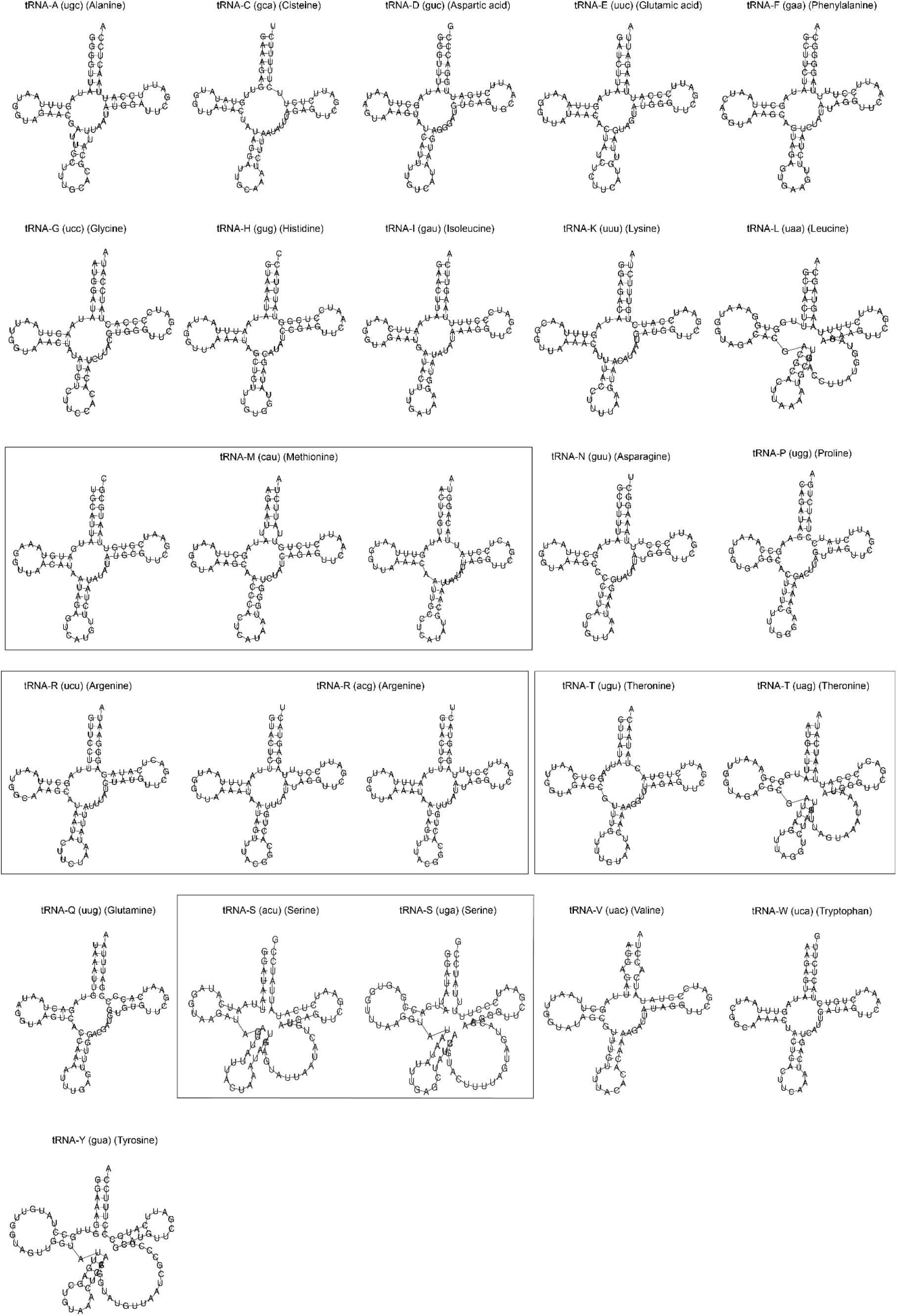
Secondary structure of 26 mt-tRNAs of *W. anomalus*.

**Figure 5.**
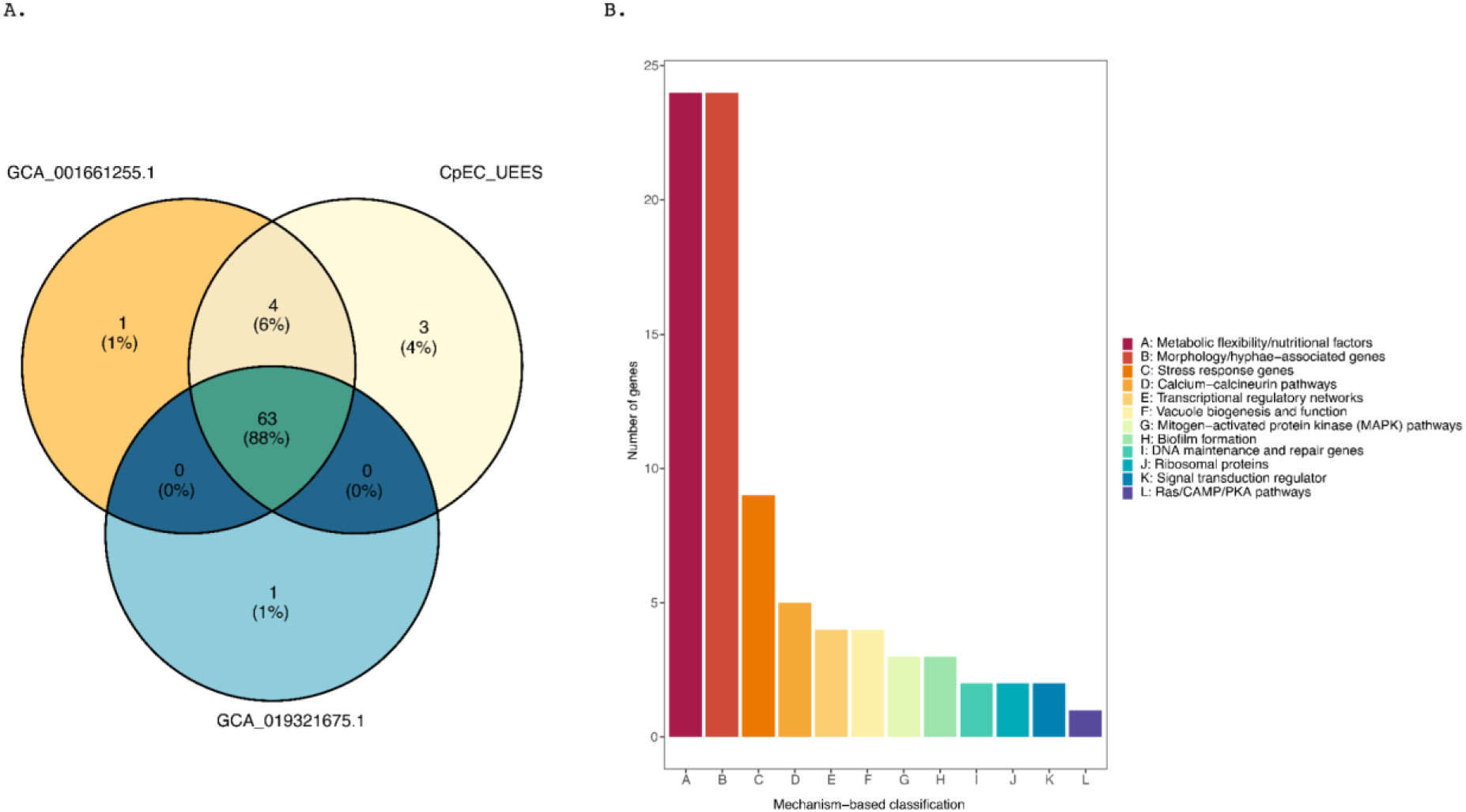
Distribution of pathogenic and virulence factors from CpEC_UEES, GCA_001661255.1, and GCA_019321675.1 genomes. A. Venn diagram showing unique and shared virulence-associated genes in CpEC_UEES and reference genomes. B. Mechanism-based classification and distribution of virulence factors found in the CpEC_UEES genome.

**Figure 6.**
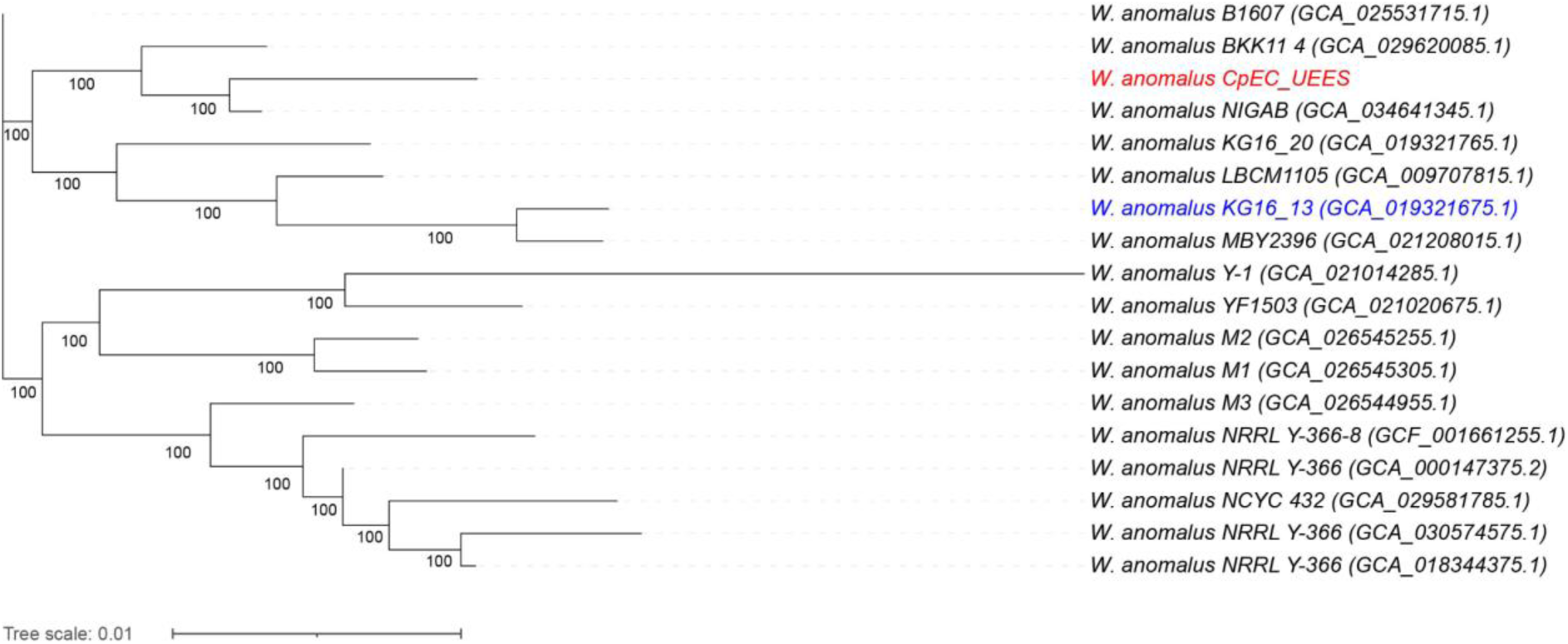
Core genome SNPs phylogeny of *W. anomalus* isolates. The unrooted phylogeny tree was inferred using IQTree with the TVM+F+I+G4 substitution model and 1,000 bootstrap replicates from an alignment of 18 core genome SNPs by SNippy, using the reference genome, and viewed on iTOL (40). Blue colour: reference genome, red colour: CpEC_UEES.

Furthermore, the mitogenome contains two rRNA genes, *rrnS* (small subunit ribosomal RNA) and *rrnL* (large subunit ribosomal RNA) with lengths of 1,822 bp and 5,083 bp, respectively. The *rrnS* was located between the *trnM* and the *trnV* genes (Figure 3), while the *rrnL* gene contains an LD-coding endonuclease (as mentioned above). While MFannot did not identify *rrnL*, initial annotation with MITOS2 predicted the gene as three separate fragments, misidentifying the region (45,192-45,272 bp) as part of an intron. This missing annotation was corrected through BLAST analysis against the *rrnL* sequences from *Wickerhamomyces mucosus* (KC993197.1) and *Saccharomyces cerevisiae* (OP414676.1 and KP712803.1), which identified three exons (Table 1). Infernal’s bacterial LSU rRNA model (RF02541) further confirmed conserved rRNA domains within exon 1 (44,838 - 44,897 bp, E-value 8.8e-6) and exon 2 (45,194 - 45,270 bp, E-value 1e-18). The final *rrnL* structure comprises three exons (41,875-44,895, 45,192-45,272, and 46,733-46,977, Table 2) separated by two intronic regions, one of which harbors the LD ORF.

### Introns, Mobile and Repetitive elements

Two group I introns hosting LD endonucleases were identified and classified into the group IA3 (positions 45,311 to 46,717, 1,407 bp) within *rrnL*, and the group ID (25,041 - 26,456, 1415 bp) within *cox1*. Additionally, RNAweasel and Infernal (group I introns model, RF00028) (36) annotated a short 218 bp region as a group IA intron (44,913–45,130) between *rrnL*’s first and second exons, further supported by exonic alignments described in the previous section.

Both MITOS2 annotation and PFAM domain signature identified an LD-homing endonuclease domain within the *cob* gene. However, neither RNAweasel nor Infernal identifies a group I/II intron structure surrounding this ORF. We then aligned the *cob* gene sequence with intronless *cob* orthologs (*Kluyveromyces lactis, Wickerhahomyces candensis,* and *Nakaseomyces glabratus*) and the intron-rich *S. cerevisiae cob* gene. The LD-HE region showed weak homology with the intronic regions bl1 and bl2 of *S. cerevisiae* (Supplementary Figure 3), but was absent in intronless species (Supplementary Figure 4). These results suggest that the LD ORF is not part of a functional group I intron, and may represent either an intron degeneration event that retained the endonuclease ORF, or an intronless homing event.

Tandem repeat analysis identified 77 repeats, ranging from 31 bp (44946-44976, 10 copies of 3 bp) to 607 bp (9615 - 10221, 1 copy of 543 bp), with 27 repeats having short units (2 −15 bp, up to 74 copies) (Supplementary table 7). Most repeats were located in intergenic regions (Supplementary Figure 5). RepeatMasker (37) identified 39 AT-rich simple sequence repeats (SSRs), with motifs such as (ATA)n, (TAA)n, and (TAT)n. Overlap analysis showed that 25 SSRs partially coincide with tandem repeats. Lastly, no transposable elements were found.

### Resistance and virulence factors

We identified a total of 70 virulence-associated proteins in the CpEC_UEES genome, 64 genes in the chromosome level genome available in the NCBI database (KG16_13) and 68 genes in the annotated reference genome (NRRL Y-366-8), which despite being an incomplete assembly, remains the only functionally annotated version (Figure 3a, Supplementary Table 8). Comparing these results with each other, we found a high number of virulence-associated genes shared among the reference genomes and the CpEC_UEES genome (63 proteins corresponding to 87.5%). We classified the determinant pathogenic and virulence factors into 11 groups based on their main mechanism and/or signalling pathways: calcium-calcineurin pathways, DNA maintenance and repair genes, metabolism flexibility/nutritional factors, mitogenic-activated protein kinase (MAPK) pathways, morphology/hyphae-associated genes, Ras/CAMP/PKA pathways, ribosomal proteins, signal transduction regulator, stress response genes, transcriptional regulatory networks, and vacuole biogenesis and function (Figure 3b). Most proteins (n=22) were involved in metabolic pathways, followed by morphology/hyphae-associated genes (n=19), and stress response genes (n=9).

Additionally, the presence of 1-3 unique genes in each genome suggests adaptations specific to individual strains. Results of the MaRDy analysis (39) yielded no matching results. This finding is consistent with the phenotypic observations, wherein we observed susceptibility to the antifungals tested.

### Prediction of adhesin function of annotated proteins

A total of 35 protein sequences were predicted as adhesins, from which 14 showed a query coverage and percentage of identity greater than 20% to known proteins from related fungal species (Table 2). Following the mechanism-based classification (see above), these sequences belong to three main groups: morphology/hyphae-associated proteins (n=8), metabolic flexibility/nutritional factors (n=2), biofilm formation (n=3), and one calcium-calcineurin pathways protein (Supplementary Table 9).

### Phylogenetic relationship

We compared all assemblies of *W. anomalus* at NCBI to the reference genome KG16 and identified the core SNPs using snippy-multi. The phylogenetic tree mainly comprises three clades that are associated with the type of strain as well as the corresponding isolation source. Genomes of isolates derived from the fermentation starters sourdough and Daqu belong to the same clade. The CpEC_UEES strain belongs to the clade composed of Brazilian, Korean, Thai, and Pakistani genomes.

In addition, most genomes have on average ∼14,000 SNPs, which correspond to a SNP frequency of approximately 0.22% suggesting that the genomes from the ecological isolates and our clinical strain are highly conserved (Figure 4).

## Discussion

Previous studies have consistently reported an increasing incidence of infections caused by uncommon agents of candidemia. Among these agents, *W. anomalus* has been classified as an emergent pathogenic yeast, which is increasingly reported as a source of infection in clinical settings, primarily affecting neonates and infants (3,21). Although it has been associated with poor clinical outcomes, therapeutic failures, and clonal outbreaks, our understanding of how its genome structure relates to its pathogenesis and virulence is limited. The present study reports the first genome sequence of a clinical isolate of *W. anomalus* from a wound secretion; the other available genomes at NCBI have an environmental origin. To reach these results, a hybrid read-based assembly approach complemented with short read polishing was chosen to combine the advantages of the Illumina short reads and PacBio long reads sequencing methods. The final draft genome consists of 20 contigs and 10 scaffolds, with an estimated genome size of 13.96Mb, 34.51% GC, and 0.03% Ns. The contig and scaffold N50 were 1 Mb and 2.1 Mb, respectively. We also identified a total of 6,324 protein-coding genes, 173 tRNAs, and 11 truncated transposable elements. Accordingly, the reference genome of *W. anomalus* presents similar genome size (14.1 Mb), GC percentage (34.5%), contig N50 (185.3 kb), scaffold N50 (2.2 Mb), and protein-coding genes (6,421). Our results show a more contiguous and higher-quality genome compared to the reference genome, despite one of the 10 scaffolds remaining unplaced. We further report on the first published complete mitochondrial genome assembly of *W. anomalus*.

### Characteristics of *W. anomalus* mitogenome

The mitochondrial genome of *W. anomalus* exhibits a total size of 47,621 bp and a GC content of 20.8%, which is consistent with previously reported yeast mitochondrial genomes (41). Core mitochondrial genes encoding components of oxidative phosphorylation (*atp6, atp8, atp9, nad1-6, nad4l, cox1-3,* and *cob*) were present. While these genes are conserved across fungal species, some fungal species may lack some core mitochondrial genes due to their migration to the nuclear genome during evolution. For instance, *Wickerhamomyces pijperi*, a close relative of *W. anomalus,* has independently lost all Complex I genes (41). No genes associated with pathogenicity were detected, in agreement with the mitochondrial genomes of non-pathogenic fungi, which focus more on fundamental metabolic activities, such as ATP production and biosynthesis, without influencing virulence or stress response. A striking feature of the *W. anomalus* mitogenome is the presence of two large inverted repeats (∼7kb each, Figure 3B). LIRs are common in chloroplast genomes, but are rare in mitochondrial genomes. In fungi, LIRs have been identified in a limited number of genera, including *Candida, Saccharomyces, Hanseniaspora, Agaricus, Termitomyces,* and *Malassezia* (42,43). These repeats function as recombination hotspots in *C. albicans* and may contribute to replication, structural rearrangements, and genome stability (44). In *Malassezia*, LIR formation has been linked to the duplication and inversion of the ancestral syntenic unit *trnR-trnL-atp9,* also found in *Ustilago maydis* and *Pseudozyma* sp (42). Interestingly, the LIRs of *W. anomalus* exhibit a similar inverted *trnR-atp9* arrangement, but lack *trnL*, suggesting that a similar recombination-driven mechanism underlies their formation. The two inverted repeats are identical to each other, indicating ongoing homologous recombination.

In fungi, the *atp9* gene is either absent from the mitochondrial genome, present as a single copy, or, in rare cases, duplicated (45) or within LIRs (42). Other key mitochondrial genes, including *atp6, nad2, nad4,* and *rrnl*, have been found within LIRs in certain fungi (42,43). Whether such inverted duplications confer a dosage effect that enhances fitness remains uncertain, given that mitochondrial gene functions are highly conserved and most fungi lack LIRs or duplicated genes in their mitogenomes. In *W. anomalus,* the presence of two *atp9* copies within the LIRs may provide a fitness advantage by ensuring stable ATP production during infection. However, further studies are needed to confirm the dosage effect or fitness advantage.

Fungal mitochondrial genomes are highly diverse, mainly due to the presence of mobile genetic elements (group I and group II introns) and repetitive sequences (41). In this study, we identified three LAGLIDADG-type HEGs, two of which are located within group I introns: one in a group ID intron in the *cox1* gene and one in a group IA3 intron in the *rrnL* gene. Most noteworthy is the presence of an LD ORF in the *cob* gene without an identifiable intronic sequence, as tools like Infernal and RNAweasel failed to detect neighboring intronic regions. This might be due to sequence degeneration reducing similarity to conserved intron models.

Notably, 48.9% (23,297 bp) of the mitogenome corresponds to non-coding content, primarily consisting of intergenic regions, intronic sequences (excluding LD ORFs), and repetitive elements. This unannotated fraction is within the typical range for yeast mitochondrial genomes in the Saccharomycetales order, such as *Saccharomyces* and *Candida*, where non-coding content can vary from 10% to over 50% (46). Of the non-coding region, 6.85% corresponds to intronic sequences excluding the LD ORFs, and RepeatMasker and equicktandem analysis revealed that 27.06% (6,304 bp) of the unannotated area consisted of repetitive elements, including tandem repeats and SSRs. These repeats, together with intron composition, may contribute to structural variation and plasticity. Whether such variation impacts the pathogenicity or adaptation of this particular strain remains uncertain, given the high diversity observed in fungal mitogenomes even at the intraspecies level (42,47). Nonetheless, mitochondrial genome variations, including intron content, have been associated with differences in virulence, pathogenicity, and drug resistance in several fungal species (19).

Given that this isolate originated from a clinical sample, it could be speculated that these mitochondrial features, including the large inverted repeats, might contribute to increased fitness in the host environment. However, this warrants further comparative and functional studies.

### SNPs Phylogeny reveals that CpEC_UEES is closely related to Brazilian, Korean, Thai, and Pakistani environmental strains

Comparative synteny analysis against the environmental strain KG16 (GCA_019321675.1) revealed that both the MaSuRCA and Canu-Smartdenovo assemblies are highly colinear with the reference genome, with only two inversions consistently detected in both. In addition, CpEC_UEES exhibited a similar number of SNPs compared to the other 17 genomes available at NCBI.

Phylogenetic tree based on SNPs revealed that CpEC_UEES and LBCM1105 strains from Ecuador and Brazil, respectively, belong to the *W. anomalus* clade of isolates from Korea, Thailand, and Pakistan, with 100% bootstrap confidence. Moreover, while genomes associated with specific sources and/or strains (e.g., daqu: GCA_021014285.1 and GCA_021020675.1; sourdough: M2 and M3; NRRL Y-366 strain: GCA_018344375.1, GCA_000147375.2, GCA_030574575.1, and GCA_001661255.1) clustered together, there is no clear evidence of geographical associations, possibly due to the limited number of available genomes.

The SNPs and phylogenetic analysis revealed the close relatedness of our clinical and other environmental isolates, suggesting that the genomic rearrangements might result from random recombination events during growth inside the patient. Our results align with other studies demonstrating that changes in the genomes of different fungal species occur more often in living hosts (48–50). However, the possibility that these genetic variations were present before infection cannot be discarded. Even further, determining whether these rearrangements play a role in virulence requires the analysis of larger sets of genomes of clinical *W. anomalus* isolates from different regions.

### Virulence-associated factors are present in clinical and ecological *Wickerhamomyces anomalus* genomes

As an opportunistic pathogen, *W. anomalus* demonstrates a capacity to colonise tissues and initiate infections in various hosts. A previous study reported its ability to induce lethal outcomes in the invertebrate model *Galleria mellonella*, highlighting its pathogenicity (51). To elucidate factors underlying its pathogenic potential, we conducted a comparative analysis between the predicted protein sequences and known virulence traits of different *Candida* species. Pathogen-host interactions revealed 72 proteins with high similarity to those in the database, of which 63 were conserved across all examined genomes. These results align with our findings that the *W. anomalus* genome exhibits substantial genomic congruence. These are further supported by Perez-Traves et al. (2021), where *W. anomalus* strains showed similar levels of pathogenicity in *G. mellonella* larvae regardless of their origin (clinical or environmental) (51).

We also identified adhesion-associated proteins that play a critical role in the infection process of pathogenic fungi. Based on the presence of a GPI-anchor domain, signal peptide sequence, and homology to known fungal adhesion proteins, we found a total of 12 unique adhesin-like GPI-anchored proteins orthologous to those present in *Candida albicans, Candida orthopsilosis*, and *Cyberlindnera fabianii*, indicating a potential predisposition to virulence and biofilm formation.

We combined and classified these determinant pathogenic and virulence factors (83 unique proteins) in 12 groups based on their main mechanisms, wherein metabolic flexibility/nutritional factors, morphology/hyphae-associated genes, and stress response genes were the most enriched groups with 68.7% (57 proteins) of the predicted protein sequences.

### *W. anomalus* contains metabolic flexibility/nutritional factors that might facilitate its adaptation to host environments

*W. anomalus* is considered a robust organism due to its ability to grow under various carbon and nitrogen sources, as well as extreme environmental stress conditions such as low pH, high osmotic pressure, and anaerobic conditions (6–8). Such remarkable metabolic versatility may play a pivotal role in the fungus’s virulence by facilitating adept nutrient assimilation and its adaptation to host microenvironments. Indeed, previous research on opportunistic fungi such as *Candida albicans, Aspergillus fumigatus,* and *Cryptococcus neoformans* highlights their metabolic flexibility as an integral part of their pathogenicity (52–54).

As expected, we identified several genes that are part of different metabolic pathways that impact *Candida* virulence. Among these, there are the glyoxylate cycle genes *ICL1* (Isocitrate lyase) and *MLS1* (Malate synthase), which are required to utilise alternative carbon sources amidst nutrient scarcity (55,56). Notably, *ICL1* is also required for prolonged survival following macrophage engulfment of *C. albicans* and *C. glabrata* (55,57).

We also found homologous sequences of the two main enzymes of the trehalose biosynthesis pathway, Tps1 (trehalose-6-phosphate synthase), and Tps2 (Trehalose-6-phosphate phosphatase), as well as the neutral trehalase enzymes Ntc1, Nth1, and Nt2. The synthesis and accumulation of the nonreducing disaccharide trehalose confer protection under stress conditions to several organisms, including *W. anomalus* (58,59). Accordingly, diminished tolerance to oxidative stress, decreased susceptibility to macrophage-mediated clearance, and lower infectivity are observed in *Candida* mutants unable to synthesise this disaccharide (60–63).

The metabolic versatility of *W. anomalus* is further demonstrated by the presence of essential transporters required for nutrient uptake and accumulation, such as Dur1_2 (urea transporter), Fth2 (iron transporter), Vht1 (biotin transporter), Ftr1 (high-affinity iron permease), and Gap1 (general amino acid permease).

### Morphology/hyphae-associated genes

Fungi exhibit morphological plasticity as a means to adapt to changing environmental conditions, including variations in nutrient availability and mating factors. The composition and organisation of the cell wall are tightly linked to the morphology of the cells, influencing their shape, size, and growth pattern (64). In this study, we identified several genes that are orthologs to the cell wall-associated genes *UTR2, PHR1*, and *PHR2*. *UTR2*, which encodes a transglycosidase that links 1,3-β-glucan to chitin, is involved in response mechanisms to cell wall-degrading enzymes (64,65). Accordingly, *Δutr2* mutants of *C. albicans* showed increased sensitivity to echinocandin treatment, altered morphology, reduced cell adhesion, and attenuated virulence in a mouse model of systemic candidiasis (64,65).

*PHR1* and *PHR2* are responsible for encoding glycosylphosphatidylinositol-anchored cell surface proteins with morphological functions (66). Although deletion of *PHR1* and *PHR2* from *C. albicans* and their respective homologous genes in *S. cerevisiae* and *C. glabrata* results in altered cell wall biogenesis, reduced adhesive properties, and morphological defects (67,68), these phenotypes are dependent on ambient pH in *C. albicans* and *C. dubliniensis* (66,69). In contrast to *Δphr1* cells, which are morphologically abnormal at alkaline pH (greater than 6), *Δphr2* mutants display morphological defects at acidic pH (70). Accordingly, the former mutants are avirulent in a mouse model of systemic candidiasis but display unaltered virulence in rat models of vaginal infection (66,71). The inverse phenotype is observed in *Δphr2* mutants.

*W. anomalus* typically grows as yeast but can adopt pseudohyphal structures (51,72). Nonetheless, pseudohyphae appeared to have a minimal influence on the virulence-associated mechanisms of cytotoxicity and damage potential (51). Adhesion, on the other hand, was proposed as a plausible mechanism of infection of this yeast, as *W. anomalus* strains showed high levels of adherence and cytotoxicity to epithelial cells (51). The role of *UTR2, PHR1*, and *PHR2* in the morphology and adhesion across evolutionarily distant fungi suggests a conservation of their functions. Thus, the homologue genes of *UTR2, PHR1,* and *PHR2* might have similar roles in the adhesion properties of *W. anomalus*; however, their impact on the virulence of the yeast should be further studied.

### Stress-response genes

The draft genome of *W. anomalus* reveals the presence of several genes that code for superoxide dismutases (SOD) and catalases (CAT). These enzymes are considered the first line of defence against oxidative stress in eukaryotic organisms. In the case of *W. anomalus*, a high activity of *SOD* genes was observed following ethanol stress (6). However, these were not sufficient to ameliorate growth inhibition and damage to cell membrane integrity. Interestingly, while studies have not explored the impact of SOD expression in *W. anomalus* under other environmental factor stresses, research on other opportunistic fungi has linked SODs to their virulence. For instance, *C. albicans Δsod1* mutants exhibited increased susceptibility to phagocytosis and superoxide radicals produced by the immune system, and attenuated virulence in a mouse model (73). Similarly, SODs protect against the oxidative burst of the host immune response in *Paracoccidioides* spp., *Histoplasma capsulatum*, and *Beauveria bassiana*, suggesting that the functions of these enzymes might be conserved (74–76).

Although shown to be inefficient under ethanol stress, *W. anomalus* SODs and CAT enzymes might be adequate to confer protection against superoxide radicals in the context of immunosenescence. This is supported by observations that senescent neutrophils present impaired oxidative response and phagocytic defects (77,78). Nonetheless, further research is required to establish the role of these enzymes in *W. anomalus* infection.

## Conclusions

Our study provides the first genome overview of a clinical isolate of *W. anomalus*, with an emphasis on the identification of homologs of virulence-associated genes in related pathogenic fungi. This report is a step forward toward the initiation of genomic studies of this robust fungus, which is responsible for several outbreaks around the world and has been associated with poor clinical outcomes. Although the isolate under investigation did not present resistance to the tested antifungals, reported resistance to flucytosine and triazoles in other isolates could pose a concern in clinical settings in the future. The mitochondrial genome of this isolate reveals distinctive features, including two LIRs and high non-coding content, which may contribute to the metabolic flexibility and industrial versatility of *W. anomalus*. Given its dual identity as both an opportunistic pathogen and an industrially valuable organism, continued genomic surveillance is essential to detect changes that may enhance its virulence or environmental adaptability.

## Methods

### 1. Clinical profile and identification of *W. anomalus*

*W. anomalus* was isolated from the wound drainage of an 84-year-old woman during a one-year surveillance in 2019-2020, where some rare non-Candida albicans (NCA) were identified for the first time in Ecuador (79). The isolate was identified using the VITEK 2 automated system, and culture strains were rearranged to 2.0 McFarland (1.8-2.2; DensiCheck, BioMérieux) supplemented with 0.45% sterile NaCl following the manufacturer’s recommendations and subsequently by sequencing of rDNA internal transcribed spacer (ITS) using primers ITS1-ITS4 (80). The antifungal susceptibility test was performed using VITEK^®^ 2 COMPACT using the AST-YS08 card (BioMérieux, Inc., United States), which measures the susceptibility against Amphotericin B, Caspofungin, Fluconazole, Micafungin, and Voriconazole. The isolate was deposited in the Mycotheque of the Laboratory of Omics Sciences, Universidad Espíritu Santo, Ecuador.

### 2. DNA extraction, Genome Library, and Genome Sequencing

The genomic DNA of *W. anomalus* was extracted using the PureLink Genomic DNA Mini Kit (Invitrogen Carlsbad, CA 92008, United States) as instructed by the manufacturer with minor modifications. Briefly, a single colony from a freshly streaked YPD broth medium into 5 ml was growing at 37 °C overnight with shaking at 200 rpm. Cell pellets were resuspended in the buffer protoplast (1M sorbitol, 1% β-mercaptoethanol, 0.25 mg of lysozyme (Sigma-Aldrich), and 0.25 mg of lyticase (Sigma-Aldrich)) and incubated at −80 ^0^C overnight. Subsequently, a digestion buffer, proteinase K, and RNase A were added. The sample was homogenized by bead beating on a Bead Ruptor (Omni International^TM^) in m/s for 5 minutes, and repeated three times. Then, the lysis was mixed with a binding buffer and ethanol, and subsequently loaded into the purification spin column. The impurities were eliminated using a wash buffer, and the DNA was eluted using the Elution buffer. The purified genomic DNA (DNAg) samples were stored at –30 °C until needed.

DNAg was then sent to Novogene Biotech Co., Ltd. (Beijing, China) for PacBio and Illumina sequencing. For long-read sequencing, HiFi SMRTbell libraries were prepared with the SMRTbell Express Template Prep Kit 2.0 (Pacific Biosciences, California, USA). The genomic DNA was fragmented, repaired, and A-tailed. SMRTbell hairpin adapters were then ligated to these prepared ends, and the AMPure PB beads (Pacific Biosciences, California, USA) were employed for the library’s concentration and purification steps. For sequencing, the library was checked for quantification and size distribution by Qubit and bioanalyzer, respectively. Finally, the library was pooled, and the sequencing process was conducted using the PacBio Sequel II platform. Reads were trimmed, and high-quality regions were conserved. A total of 3.03 Gb was obtained, with a N50 of 173.928 bp.

For short-reads, the libraries and quality control were generated using a total amount of 1.0μg DNA. Sequencing libraries were generated using NEBNext® DNA Library Prep Kit following the manufacturer’s recommendations. The genomic DNA was randomly fragmented to a size of 350 bp by shearing, then the DNA fragments were end polished, A-tailed, and ligated with the NEBNext adapter for Illumina sequencing, and further PCR enriched by P5 and indexed P7 oligos. The PCR products were purified (AMPure XP system), and the resulting libraries were analysed for size distribution by Agilent 2100 Bioanalyzer and quantified using real-time PCR. For sequencing, the qualified libraries are fed into Illumina sequencers after pooling according to their effective concentration and expected data volume. The DNA was sequenced using the Illumina NovaSeq 6000 platform, generating 9.1 Gb paired-end reads (150 bp each) with 93.38% Q30. Read quality was assessed using FastQC, and adapter sequences and low-quality bases were removed by quality-filtering and trimming with Trimmomatic (version 0.39).

### 3. Genome Assembly

Highly accurate single-molecule consensus (CCS) reads were obtained by alignment of the subreads using the pbccs tool. Then, the CCS reads were used for genome assembly using different strategies. The first approach used only the generated long reads, using two different assemblers, SMARTdenovo (28) and Flye (27). A second approach consisted of using the hybrid assemblers MaSuRCA (26) and Wengan (25) with default parameters. A fifth genome assembly, denominated as CANU-SMARTdenovo assembly, was built by adding a correction step of long reads using CANU (81) using the LRSDAY (82) pipeline.

The set of contigs produced by each assembly strategy were corrected three consecutive times with the Pilon software (83) using the quality-filtered Illumina paired-end reads. The most contiguous assembly was then scaffolded with RagTag (29), using the reference genome GCA_019321675.1.

#### 3.1 Assembly assessment

Several quality metrics were collected to assess the quality of the genome assembly. Assembly contiguity was assessed using QUAST (84) and included a total number of contigs, total sequence, and contig NG50, among others. Reference-based analysis was performed using the published reference genome GCA_019321675.1.

The completeness of the assembly was measured using the BUSCO tool with the Saccharomycetes database. Key assembly metrics, including N50, L50, longest contigs, number of contigs, GC content, and BUSCO scores, have been represented as SnailPlots using BlobToolKit (31).

Assembly QV was determined using Meryl and Merqury (85). The Meryl databases were generated for each read file (PacBio long reads and Illumina paired-end reads) with a k-mer size of 17. Merqury was then run based on each Meryl database to determine the assembly QV.

### 4. Genome Annotation

The genome was annotated using an adapted MAKER-based pipeline in the LRSDAY pipeline (82,86). Two different *ab initio* gene predictors were used: SNAP v2013_11_29 (87) and Augustus (88,89), with soft masking in the masking repeating step. Augustus was trained using a *S. cerevisiae* model. Additionally, available protein sequences of *W. anomalus, C. glabrata, S. cerevisiae* (strain S288c), and *C. kefyr* were provided as homology evidence. A similar approach was performed to annotate the reference genomes: GCA_019321675.1 and GCA_001661255.1. tRNA genes were annotated using tRNAscan-SE (90), and mobile elements were predicted based on the predetermined TE-library available at LRSDAY. Annotation quality and completeness were assessed by BUSCO (30). Functional annotation was performed by mapping the predicted protein sequences to annotated orthologous groups using eggNOG-mapper v2 (33,91) and BlastKOALA (92).

### 5. Mitochondrial Identification

The mitochondrial genome of CpEc_Uees was identified based on strong protein homology to the mitochondrial genes of *Saccharomyces cerevisiae* (strain S288c), *C. kefyr* (strain DMKU3-1042), and *C. glabrata* (strain ATCC 2001). This contig, along with contigs scf7180000000191 and scf7180000000194, were used as a template onto which the PacBio CCS reads were mapped using minimap2 v2.26 (93). We then performed an independent assembly of the mapped long reads using CANU, which resulted in a circularizable contig corresponding to the mitogenome and another collapsed contig. The assembly was then corrected using Illumina short reads (as described in Section 3).

MitoHifi (94) was used to identify the mitochondrial sequence and circularize it. This was followed by annotation with MITOS2 (34) and MFannot (35), and intron identification using RNAweasel (35) and Infernal v1.1.5 (36). Annotation inconsistencies were manually verified using BLAST. Repeats were identified using RepeatMasker (37) and equicktandem (95).

### 6. Identification of genes related to virulence/drug resistance

The PHI-base (Pathogen-host-interactions database) (38) and MaRDy (Mycology Antifungal Resistance Database) (39) databases were used for the identification of virulence and resistance genes, respectively. A BLASTP analysis was performed with an E-value cut-off of 1e-6 using the predicted protein sequences as queries against the databases. Only hits from *Candida spp.* and with a percent identity and query coverage greater than 60% and 80%, respectively, were considered. For further confirmation, the hits were compared against the genes identified by the eggNOG-mapper (33,91).

### 7. Identification of adhesin-like protein sequences

We used multiple strategies to identify adhesin-related sequence structures. First, a preliminary prediction for a signal peptide was performed with the SignalP 6.0 server (96). The organism group was set to Eukarya. We then combined these results with predictions for a C-terminal Glycosylphosphatidylinositol (GPI) anchor sequence. GPI-anchored proteins were identified using the PredGPI predictor (97). Lastly, these sequences were input into FungalRV (98), a Machine learning fungal adhesion predictor trained with known adhesin protein sequences from 8 human fungal pathogens. Proteins with a Factor score greater than zero were considered positive hits. Proteins that passed these filters were subjected to blastp against the non-redundant protein database limited to *Candida (*taxid: 5475), *Wickerhamomyces* (taxid: 599737), and *Cyberlindnera fabianii* (taxid: 36022). When no results were obtained, a general blastp was performed.

### 8. Phylogenetics Analysis

The snippy-multi protocol was used to analyse the phylogenetic relationship between the CpEC_Uees strain and other environmental *W. anomalus* strains retrieved from the NCBI database based on core SNP differences. The alignment of Core genome SNPs was produced using snippy-multi in Snippy V4.6.0 (99). Available from: https://github.com/tseemann/snippy.

The phylogenetic tree was constructed using IQ-TREE (100) with the TVM+F+I+G4 substitution model, selected by ModelFinder (101), and 1,000 bootstrap replicates. The figure was visualised using Interactive Tree of Life (iTOL) (40).

### 9. Data visualisation

Figures were constructed in R version 4.2.2 for Mac using the packages ggplot2 (102) and ggbreak (103). Additionally, pyGenomeviz and pyCirclize were used to visualise genome alignments and the mitogenome.

## Abbreviations

BUSCO: Benchmarking Universal Single-Copy Orthologs
GO: Gene Ontology
KOG: euKaryotic Orthologous Groups
KEGG: Kyoto Encyclopedia of Genes and Genomes
RSCU: Relative Synonymous Codon Usage
ORF: Open Reading Frame (marco de lectura abierto)
SSR: Simple Sequence Repeat (repeticiones simples)
SNPs: Single-Nucleotide Polymorphism
GPI: Glycosylphosphatidylinositol
PCG: Protein-Coding Gene (gen codificante de proteína)
tRNA: Transfer RNA
rRNA: Ribosomal RNA
rrnS: small subunit ribosomal RNA (15S)
rrnL: large subunit ribosomal RNA (21S)
HE: Homing Endonuclease
ccs: Circular Consensus Sequencing
MIC: Minimum Inhibitory Concentration

## Acknowledgements

Acknowledge to: GAR scholarships, SENESCYT Scholarship Number AR5G-000125-2016

## Funding

We acknowledge the support from Centro de Investigaciones, Universidad Espiritu Santo

## Availability of data and materials

Whole Genome sequencing data have been deposited in the NCBI database under the BioSample accession number SAMN39985750.

## Author information

### Authors Contribution

MCV: Perform data analysis, and draft the manuscript, LSCS and ACV: Perform data analysis, YA, JCFC and GML: Perform experiments, JU: Draft revision, DAM: Conceived the study, supervised the research, and revised the manuscript. All authors have read and approved the manuscript.

## Ethics declarations

This study was approved by the Ethics Committee of Universidad Espiritu Santo (CEISH-UEES), project Number 2022-001A. All data was anonymized.

## Consent for publication

Not applicable.

## Competing interests

The authors declare that they have no competing interests.

## Publisher’s Note

Springer Nature remains neutral with regard to jurisdictional claims in published maps and institutional affiliations

## References

1. Soriano A, Honore PM, Puerta-Alcalde P, Garcia-Vidal C, Pagotto A, Gonçalves-Bradley DC, et al. Invasive candidiasis: current clinical challenges and unmet needs in adult populations. Journal of Antimicrobial Chemotherapy. 2023 Jul 1;78(7):1569–85.

2. Rokas A. Evolution of the human pathogenic lifestyle in fungi. Nat Microbiol. 2022 May;7(5):607–19.

3. Sharma M, Chakrabarti A. Candidiasis and Other Emerging Yeasts. Curr Fungal Infect Rep. 2023 Mar 1;17(1):15–24.

4. Rayens E, Norris KA. Prevalence and Healthcare Burden of Fungal Infections in the United States, 2018. Open Forum Infectious Diseases. 2022 Jan 1;9(1):ofab593.

5. Ioannou P, Baliou S, Kofteridis DP. Fungemia by Wickerhamomyces anomalus—A Narrative Review. Pathogens. 2024 Mar 21;13(3):269.

6. Li Y, Long H, Jiang G, Gong X, Yu Z, Huang M, et al. Analysis of the ethanol stress response mechanism in Wickerhamomyces anomalus based on transcriptomics and metabolomics approaches. BMC Microbiology. 2022 Nov 15;22(1):275.

7. da Cunha AC, Gomes LS, Godoy-Santos F, Faria-Oliveira F, Teixeira JA, Sampaio GMS, et al. High-affinity transport, cyanide-resistant respiration, and ethanol production under aerobiosis underlying efficient high glycerol consumption by Wickerhamomyces anomalus. Journal of Industrial Microbiology and Biotechnology. 2019 May 1;46(5):709–23.

8. Passoth V, Fredlund E, Druvefors UÄ, Schnürer J. Biotechnology, physiology and genetics of the yeast Pichia anomala. FEMS Yeast Research. 2006 Jan 1;6(1):3–13.

9. Padilla B, Gil JV, Manzanares P. Challenges of the Non-Conventional Yeast Wickerhamomyces anomalus in Winemaking. Fermentation. 2018 Sep;4(3):68.

10. Cappelli A, Favia G, Ricci I. Wickerhamomyces anomalus in Mosquitoes: A Promising Yeast-Based Tool for the “Symbiotic Control” of Mosquito-Borne Diseases. Front Microbiol. 2021 Jan 21;11:621605.

11. Cecarini V, Cuccioloni M, Bonfili L, Ricciutelli M, Valzano M, Cappelli A, et al. Identification of a Killer Toxin from Wickerhamomyces anomalus with β-Glucanase Activity. Toxins. 2019 Sep 28;11(10):568.

12. Brown AJP, Brown GD, Netea MG, Gow NAR. Metabolism impacts upon Candida immunogenicity and pathogenicity at multiple levels. Trends in Microbiology. 2014 Nov;22(11):614–22.

13. Calderone R, Li D, Traven A. System-level impact of mitochondria on fungal virulence: to metabolism and beyond. FEMS Yeast Research [Internet]. 2015 Jun 1 [cited 2024 Sep 11];15(4). Available from: http://academic.oup.com/femsyr/article/doi/10.1093/femsyr/fov027/645474/Systemlevel-impact-of-mitochondria-on-fungal

14. Sun N, Parrish RS, Calderone RA, Fonzi WA. Unique, Diverged, and Conserved Mitochondrial Functions Influencing Candida albicans Respiration. Ljungdahl PO, Hube B, editors. mBio. 2019 Jun 25;10(3):e00300–19.

15. Misas E, Chow NA, Gómez OM, Muñoz JF, McEwen JG, Litvintseva AP, et al. Mitochondrial Genome Sequences of the Emerging Fungal Pathogen Candida auris. Front Microbiol. 2020 Oct 27;11:560332.

16. Limits to Sequencing and de novo Assembly: Classic Benchmark Sequences for Optimizing Fungal NGS Designs. In: Advances in Intelligent Systems and Computing [Internet]. Cham: Springer International Publishing; 2014 [cited 2025 Jul 23]. p. 221–30. Available from: https://link.springer.com/10.1007/978-3-319-01568-2_32

17. Li Q, Wang Q, Jin X, Chen Z, Xiong C, Li P, et al. The first complete mitochondrial genome from the family Hygrophoraceae (Hygrophorus russula) by next-generation sequencing and phylogenetic implications. International Journal of Biological Macromolecules. 2019 Feb;122:1313–20.

18. Song N, Geng Y, Li X. The Mitochondrial Genome of the Phytopathogenic Fungus Bipolaris sorokiniana and the Utility of Mitochondrial Genome to Infer Phylogeny of Dothideomycetes. Front Microbiol [Internet]. 2020 May 8 [cited 2025 Jul 23];11. Available from: https://www.frontiersin.org/journals/microbiology/articles/10.3389/fmicb.2020.00863/full

19. Ni Y, Gao X. Uncovering the role of mitochondrial genome in pathogenicity and drug resistance in pathogenic fungi. Front Cell Infect Microbiol. 2025 Apr 16;15:1576485.

20. Spruijtenburg B, Rudramurthy SM, Meijer EFJ, Van Haren MHI, Kaur H, Chakrabarti A, et al. Application of Novel Short Tandem Repeat Typing for Wickerhamomyces anomalus Reveals Simultaneous Outbreaks within a Single Hospital. Microorganisms. 2023 Jun 8;11(6):1525.

21. Paul P, Pandey N, Gupta M, Kumar D, Tilak R. A Case series of Wickerhamomyces anomalus: An emerging fungal pathogen and an entity of concern for neonatal intensive care unit. Int J Med Rev Case Rep. 2020; (Reports in Neurology, Neurosur):1.

22. Dutra VR, Silva LF, Oliveira ANM, Beirigo EF, Arthur VM, Bernardes da Silva R, et al. Fatal Case of Fungemia by Wickerhamomyces anomalus in a Pediatric Patient Diagnosed in a Teaching Hospital from Brazil. JoF. 2020 Aug 25;6(3):147.

23. Zhang L, Xiao M, Arastehfar A, Ilkit M, Zou J, Deng Y, et al. Investigation of the Emerging Nosocomial Wickerhamomyces anomalus Infections at a Chinese Tertiary Teaching Hospital and a Systemic Review: Clinical Manifestations, Risk Factors, Treatment, Outcomes, and Anti-fungal Susceptibility. Front Microbiol. 2021 Oct 6;12:744502.

24. Barchiesi F, Tortorano AM, Di Francesco LF, Rigoni A, Giacometti A, Spreghini E, et al. Genotypic variation and antifungal susceptibilities of Candida pelliculosa clinical isolates. Journal of Medical Microbiology. 2005 Mar 1;54(3):279–85.

25. Di Genova A, Buena-Atienza E, Ossowski S, Sagot MF. Efficient hybrid de novo assembly of human genomes with WENGAN. Nat Biotechnol. 2021 Apr;39(4):422–30.

26. Zimin AV, Puiu D, Luo MC, Zhu T, Koren S, Marçais G, et al. Hybrid assembly of the large and highly repetitive genome of Aegilops tauschii, a progenitor of bread wheat, with the MaSuRCA mega-reads algorithm. Genome Res. 2017 May;27(5):787–92.

27. Kolmogorov M, Yuan J, Lin Y, Pevzner PA. Assembly of long, error-prone reads using repeat graphs. Nat Biotechnol. 2019 May;37(5):540–6.

28. Liu H, Wu S, Li A, Ruan J. SMARTdenovo: a de novo assembler using long noisy reads. Gigabyte. 2021 Mar 8;2021:1–9.

29. Alonge M, Lebeigle L, Kirsche M, Jenike K, Ou S, Aganezov S, et al. Automated assembly scaffolding using RagTag elevates a new tomato system for high-throughput genome editing. Genome Biology. 2022 Dec 15;23(1):258.

30. Manni M, Berkeley MR, Seppey M, Simão FA, Zdobnov EM. BUSCO Update: Novel and Streamlined Workflows along with Broader and Deeper Phylogenetic Coverage for Scoring of Eukaryotic, Prokaryotic, and Viral Genomes. Molecular Biology and Evolution. 2021 Oct 1;38(10):4647–54.

31. Challis R, Richards E, Rajan J, Cochrane G, Blaxter M. BlobToolKit – Interactive Quality Assessment of Genome Assemblies. G3 Genes|Genomes|Genetics. 2020 Apr 1;10(4):1361–74.

32. Marçais G, Delcher AL, Phillippy AM, Coston R, Salzberg SL, Zimin A. MUMmer4: A fast and versatile genome alignment system. PLOS Computational Biology. 2018 Jan 26;14(1):e1005944.

33. Cantalapiedra CP, Hernández-Plaza A, Letunic I, Bork P, Huerta-Cepas J. eggNOG-mapper v2: Functional Annotation, Orthology Assignments, and Domain Prediction at the Metagenomic Scale. Molecular Biology and Evolution. 2021 Dec 1;38(12):5825–9.

34. Bernt M, Donath A, Jühling F, Externbrink F, Florentz C, Fritzsch G, et al. MITOS: Improved *de novo* metazoan mitochondrial genome annotation. Molecular Phylogenetics and Evolution. 2013 Nov 1;69(2):313–9.

35. Lang BF, Beck N, Prince S, Sarrasin M, Rioux P, Burger G. Mitochondrial genome annotation with MFannot: a critical analysis of gene identification and gene model prediction. Front Plant Sci [Internet]. 2023 Jul 4 [cited 2025 Jul 23];14. Available from: https://www.frontiersin.org/journals/plant-science/articles/10.3389/fpls.2023.1222186/full

36. Nawrocki EP, Eddy SR. Infernal 1.1: 100-fold faster RNA homology searches. Bioinformatics. 2013 Nov 15;29(22):2933–5.

37. Smit, AFA, Hubley, R, Green, P. RepeatMasker Open-4.0 [Internet]. 2013. Available from: http://www.repeatmasker.org

38. Urban M, Cuzick A, Seager J, Wood V, Rutherford K, Venkatesh SY, et al. PHI-base in 2022: a multi-species phenotype database for Pathogen–Host Interactions. Nucleic Acids Research. 2022 Jan 7;50(D1):D837–47.

39. Nash A, Sewell T, Farrer RA, Abdolrasouli A, Shelton JMG, Fisher MC, et al. MARDy: Mycology Antifungal Resistance Database. Kelso J, editor. Bioinformatics. 2018 Sep 15;34(18):3233–4.

40. Letunic I, Bork P. Interactive tree of life (iTOL) v3: an online tool for the display and annotation of phylogenetic and other trees. Nucleic Acids Res. 2016 Jul 8;44(W1):W242–5.

41. Wolters JF, LaBella AL, Opulente DA, Rokas A, Hittinger CT. Mitochondrial genome diversity across the subphylum Saccharomycotina. Front Microbiol. 2023 Nov 23;14:1268944.

42. Christinaki AC, Theelen B, Zania A, Coutinho SDA, Cabañes JF, Boekhout T, et al. Co-evolution of large inverted repeats and G-quadruplex DNA in fungal mitochondria may facilitate mitogenome stability: the case of Malassezia. Sci Rep. 2023 Apr 18;13(1):6308.

43. Nieuwenhuis M, van de Peppel LJJ, Bakker FT, Zwaan BJ, Aanen DK. Enrichment of G4DNA and a Large Inverted Repeat Coincide in the Mitochondrial Genomes of Termitomyces. Genome Biol Evol. 2019 Jul 1;11(7):1857–69.

44. Gerhold JM, Aun A, Sedman T, Jõers P, Sedman J. Strand Invasion Structures in the Inverted Repeat of *Candida albicans* Mitochondrial DNA Reveal a Role for Homologous Recombination in Replication. Molecular Cell. 2010 Sep 24;39(6):851–61.

45. Duò A, Bruggmann R, Zoller S, Bernt M, Grünig CR. Mitochondrial genome evolution in species belonging to the Phialocephala fortinii s.l. - Acephala applanata species complex. BMC Genomics. 2012 May 4;13(1):166.

46. Medina R, Franco MEE, Bartel LC, Martinez Alcántara V, Saparrat MCN, Balatti PA. Fungal Mitogenomes: Relevant Features to Planning Plant Disease Management. Front Microbiol [Internet]. 2020 May 29 [cited 2025 Jul 18];11. Available from: https://www.frontiersin.org/journals/microbiology/articles/10.3389/fmicb.2020.00978/full

47. Christinaki AC, Kanellopoulos SG, Kortsinoglou AM, Andrikopoulos MΑ, Theelen B, Boekhout T, et al. Mitogenomics and mitochondrial gene phylogeny decipher the evolution of *Saccharomycotina* yeasts. Wolfe K, editor. Genome Biology and Evolution. 2022 May 3;14(5):evac073.

48. Forche A, Magee PT, Selmecki A, Berman J, May G. Evolution in Candida albicans Populations During a Single Passage Through a Mouse Host. Genetics. 2009 Jul;182(3):799–811.

49. Smith AC, Hickman MA. Host-Induced Genome Instability Rapidly Generates Phenotypic Variation across Candida albicans Strains and Ploidy States. Mitchell AP, editor. mSphere. 2020 Jun 24;5(3):e00433–20.

50. Grimberg B, Zeyl C. The Effects Of Sex And Mutation Rate On Adaptation In Test Tubes And To Mouse Hosts By Saccharomyces Cerevisiae. Evolution. 2005 Feb;59(2):431–8.

51. Peréz-Través L, de Llanos R, Flockhart A, García-Domingo L, Groenewald M, Pérez-Torrado R, et al. Virulence related traits in yeast species associated with food; Debaryomyces hansenii, Kluyveromyces marxianus, and Wickerhamomyces anomalus. Food Control. 2021 Jun 1;124:107901.

52. Garbe E, Vylkova S. Role of Amino Acid Metabolism in the Virulence of Human Pathogenic Fungi. Curr Clin Micro Rpt. 2019 Sep 1;6(3):108–19.

53. Fourie R, Kuloyo OO, Mochochoko BM, Albertyn J, Pohl CH. Iron at the Centre of Candida albicans Interactions. Front Cell Infect Microbiol. 2018 Jun 5;8:185.

54. Amich J, Krappmann S. Deciphering metabolic traits of the fungal pathogen Aspergillus fumigatus: redundancy vs. essentiality. Frontiers in Microbiology [Internet]. 2012 [cited 2024 Feb 14];3. Available from: https://www.frontiersin.org/journals/microbiology/articles/10.3389/fmicb.2012.00414

55. Chew SY, Ho KL, Cheah YK, Ng TS, Sandai D, Brown AJP, et al. Glyoxylate cycle gene ICL1 is essential for the metabolic flexibility and virulence of Candida glabrata. Sci Rep. 2019 Feb 26;9(1):2843.

56. Lorenz MC, Fink GR. The glyoxylate cycle is required for fungal virulence. Nature. 2001 Jul;412(6842):83–6.

57. Ramírez MA, Lorenz MC. Mutations in Alternative Carbon Utilization Pathways in Candida albicans Attenuate Virulence and Confer Pleiotropic Phenotypes. Eukaryot Cell. 2007 Feb;6(2):280–90.

58. Fredlund E, Broberg A, Boysen ME, Kenne L, Schnürer J. Metabolite profiles of the biocontrol yeast Pichia anomala J121 grown under oxygen limitation. Appl Microbiol Biotechnol. 2004 Apr 1;64(3):403–9.

59. Chen Q, Haddad GG. Role of trehalose phosphate synthase and trehalose during hypoxia: from flies to mammals. Journal of Experimental Biology. 2004 Aug 15;207(18):3125–9.

60. Martínez-Esparza M, Tapia-Abellán A, Vitse-Standaert A, García-Peñarrubia P, Argüelles JC, Poulain D, et al. Glycoconjugate expression on the cell wall of tps1/tps1 trehalose-deficient Candida albicans strain and implications for its interaction with macrophages. Glycobiology. 2011 Jun 1;21(6):796–805.

61. Zaragoza O, Blazquez MA, Gancedo C. Disruption of the Candida albicans TPS1Gene Encoding Trehalose-6-Phosphate Synthase Impairs Formation of Hyphae and Decreases Infectivity. Journal of Bacteriology. 1998 Aug;180(15):3809–15.

62. Van Dijck P, De Rop L, Szlufcik K, Van Ael E, Thevelein JM. Disruption of the Candida albicans TPS2 Gene Encoding Trehalose-6-Phosphate Phosphatase Decreases Infectivity without Affecting Hypha Formation. Infection and Immunity. 2002 Apr;70(4):1772–82.

63. Van Ende M, Timmermans B, Vanreppelen G, Siscar-Lewin S, Fischer D, Wijnants S, et al. The involvement of the Candida glabrata trehalase enzymes in stress resistance and gut colonization. Virulence. 2021 Dec 31;12(1):329–45.

64. Pardini G, Groot PWJD, Coste AT, Karababa M, Klis FM, Koster CG de, et al. The CRH Family Coding for Cell Wall Glycosylphosphatidylinositol Proteins with a Predicted Transglycosidase Domain Affects Cell Wall Organization and Virulence of Candida albicans*. Journal of Biological Chemistry. 2006 Dec 29;281(52):40399–411.

65. Walker LA, Munro CA. Caspofungin Induced Cell Wall Changes of Candida Species Influences Macrophage Interactions. Front Cell Infect Microbiol. 2020 May 12;10:164.

66. Calderon J, Zavrel M, Ragni E, Fonzi WA, Rupp S, Popolo L. PHR1, a pH-regulated gene of Candida albicans encoding a glucan-remodelling enzyme, is required for adhesion and invasion. Microbiology. 2010;156(8):2484–94.

67. Popolo L, Vai M, Gatti E, Porello S, Bonfante P, Balestrini R, et al. Physiological analysis of mutants indicates involvement of the Saccharomyces cerevisiae GPI-anchored protein gp115 in morphogenesis and cell separation. J Bacteriol. 1993 Apr;175(7):1879–85.

68. Weig M, Haynes K, Rogers TR, Kurzai O, Frosch M, Mühlschlegel FA. A GAS-like gene family in the pathogenic fungus Candida glabrataThe EMBL accession numbers for the sequences reported in this paper are AJ302061 for CgGAS1, AJ302062 for CgGAS2 and AJ302063 for CgGAS3. Microbiology. 2001;147(8):2007–19.

69. Heinz WJ, Kurzai O, Brakhage AA, Fonzi WA, Korting HC, Frosch M, et al. Molecular responses to changes in the environmental pH are conserved between the fungal pathogens Candida dubliniensis and Candida albicans. International Journal of Medical Microbiology. 2000 Jul;290(3):231–8.

70. Mühlschlegel FA, Fonzi WA. PHR2 of Candida albicans encodes a functional homolog of the pH-regulated gene PHR1 with an inverted pattern of pH-dependent expression. Mol Cell Biol. 1997 Oct;17(10):5960–7.

71. De Bernardis F, Mühlschlegel FA, Cassone A, Fonzi WA. The pH of the Host Niche Controls Gene Expression in and Virulence of Candida albicans. Infect Immun. 1998 Jul;66(7):3317–25.

72. Kurtzman CP. *Pichia* E.C. Hansen emend. Kurtzman. In: Kurtzman CP, Fell JW, editors. The Yeasts (Fourth Edition) [Internet]. Amsterdam: Elsevier; 1998 [cited 2024 Feb 16]. p. 273–352. Available from: https://www.sciencedirect.com/science/article/pii/B9780444813121500460

73. Hwang CS, Rhie G eun, Oh JH, Huh WK, Yim HS, Kang SO. Copper- and zinc-containing superoxide dismutase (Cu/ZnSOD) is required for the protection of Candida albicans against oxidative stresses and the expression of its full virulence. Microbiology. 2002;148(11):3705–13.

74. Youseff BH, Holbrook ED, Smolnycki KA, Rappleye CA. Extracellular Superoxide Dismutase Protects Histoplasma Yeast Cells from Host-Derived Oxidative Stress. PLOS Pathogens. 2012 May 17;8(5):e1002713.

75. Tamayo D, Muñoz JF, Lopez Á, Urán M, Herrera J, Borges CL, et al. Identification and Analysis of the Role of Superoxide Dismutases Isoforms in the Pathogenesis of Paracoccidioides spp. PLOS Neglected Tropical Diseases. 2016 Mar 10;10(3):e0004481.

76. Li F, Shi HQ, Ying SH, Feng MG. Distinct contributions of one Fe- and two Cu/Zn-cofactored superoxide dismutases to antioxidation, UV tolerance and virulence of *Beauveria bassiana*. Fungal Genetics and Biology. 2015 Aug 1;81:160–71.

77. Moss CE, Phipps H, Wilson HL, Kiss-Toth E. Markers of the ageing macrophage: a systematic review and meta-analysis. Frontiers in Immunology [Internet]. 2023 [cited 2024 Feb 18];14. Available from: https://www.frontiersin.org/journals/immunology/articles/10.3389/fimmu.2023.1222308

78. Andrade B, Jara-Gutiérrez C, Paz-Araos M, Vázquez MC, Díaz P, Murgas P. The Relationship between Reactive Oxygen Species and the cGAS/STING Signaling Pathway in the Inflammaging Process. International Journal of Molecular Sciences. 2022 Jan;23(23):15182.

79. Acosta-Mosquera Y, Tapia JC, Armas-González R, Cáceres-Valdiviezo MJ, Fernández-Cadena JC, Andrade-Molina D. Prevalence and Species Distribution of Candida Clinical Isolates in a Tertiary Care Hospital in Ecuador Tested from January 2019 to February 2020. JoF. 2024 Apr 24;10(5):304.

80. White TJ, Bruns T, Lee S, Taylor J. Amplification And Direct Sequencing Of Fungal Ribosomal Rna Genes For Phylogenetics. In: PCR Protocols [Internet]. Elsevier; 1990 [cited 2024 Jan 29]. p. 315–22. Available from: https://linkinghub.elsevier.com/retrieve/pii/B9780123721808500421

81. Koren S, Walenz BP, Berlin K, Miller JR, Bergman NH, Phillippy AM. Canu: scalable and accurate long-read assembly via adaptive k-mer weighting and repeat separation. Genome Res. 2017 May 1;27(5):722–36.

82. Yue JX, Liti G. Long-read sequencing data analysis for yeasts. Nat Protoc. 2018 Jun;13(6):1213–31.

83. Walker BJ, Abeel T, Shea T, Priest M, Abouelliel A, Sakthikumar S, et al. Pilon: An Integrated Tool for Comprehensive Microbial Variant Detection and Genome Assembly Improvement. PLOS ONE. 2014 Nov 19;9(11):e112963.

84. Gurevich A, Saveliev V, Vyahhi N, Tesler G. QUAST: quality assessment tool for genome assemblies. Bioinformatics. 2013 Apr 15;29(8):1072–5.

85. Rhie A, Walenz BP, Koren S, Phillippy AM. Merqury: reference-free quality, completeness, and phasing assessment for genome assemblies. Genome Biol. 2020 Dec;21(1):245.

86. Holt C, Yandell M. MAKER2: an annotation pipeline and genome-database management tool for second-generation genome projects. BMC Bioinformatics. 2011 Dec 22;12:491.

87. Korf I. Gene finding in novel genomes. BMC Bioinformatics. 2004 May 14;5(1):59.

88. Stanke M, Keller O, Gunduz I, Hayes A, Waack S, Morgenstern B. AUGUSTUS: ab initio prediction of alternative transcripts. Nucleic Acids Research. 2006 Jul 1;34(suppl_2):W435–9.

89. Stanke M, Diekhans M, Baertsch R, Haussler D. Using native and syntenically mapped cDNA alignments to improve de novo gene finding. Bioinformatics. 2008 Mar 1;24(5):637–44.

90. Chan PP, Lin BY, Mak AJ, Lowe TM. tRNAscan-SE 2.0: improved detection and functional classification of transfer RNA genes. Nucleic Acids Research. 2021 Sep 20;49(16):9077–96.

91. Huerta-Cepas J, Szklarczyk D, Heller D, Hernández-Plaza A, Forslund SK, Cook H, et al. eggNOG 5.0: a hierarchical, functionally and phylogenetically annotated orthology resource based on 5090 organisms and 2502 viruses. Nucleic Acids Research. 2019 Jan 8;47(D1):D309–14.

92. Kanehisa M, Sato Y, Morishima K. BlastKOALA and GhostKOALA: KEGG Tools for Functional Characterization of Genome and Metagenome Sequences. Journal of Molecular Biology. 2016 Feb;428(4):726–31.

93. Li H. Minimap2: pairwise alignment for nucleotide sequences. Birol I, editor. Bioinformatics. 2018 Sep 15;34(18):3094–100.

94. Uliano-Silva M, Ferreira JGRN, Krasheninnikova K, Darwin Tree of Life Consortium, Blaxter M, Mieszkowska N, et al. MitoHiFi: a python pipeline for mitochondrial genome assembly from PacBio high fidelity reads. BMC Bioinformatics [Internet]. 2023 Jul 18 [cited 2025 Jul 23];24(1). Available from: https://bmcbioinformatics.biomedcentral.com/articles/10.1186/s12859-023-05385-y

95. Rice P, Longden I, Bleasby A. EMBOSS: The European Molecular Biology Open Software Suite. Trends in Genetics. 2000 Jun 1;16(6):276–7.

96. Teufel F, Almagro Armenteros JJ, Johansen AR, Gíslason MH, Pihl SI, Tsirigos KD, et al. SignalP 6.0 predicts all five types of signal peptides using protein language models. Nat Biotechnol. 2022 Jul;40(7):1023–5.

97. Pierleoni A, Martelli PL, Casadio R. PredGPI: a GPI-anchor predictor. BMC Bioinformatics. 2008 Dec;9(1):392.

98. Chaudhuri R, Ansari FA, Raghunandanan MV, Ramachandran S. FungalRV: adhesin prediction and immunoinformatics portal for human fungal pathogens. BMC Genomics. 2011 Apr 15;12:192.

99. Torsten Seemann. Snippy: rapid haploid variant calling and core SNP phylogeny [Internet]. 2015. Available from: GitHub. Available at: github. com/tseemann/snippy

100. Nguyen LT, Schmidt HA, Von Haeseler A, Minh BQ. IQ-TREE: A Fast and Effective Stochastic Algorithm for Estimating Maximum-Likelihood Phylogenies. Molecular Biology and Evolution. 2015 Jan;32(1):268–74.

101. Kalyaanamoorthy S, Minh BQ, Wong TKF, Von Haeseler A, Jermiin LS. ModelFinder: fast model selection for accurate phylogenetic estimates. Nat Methods. 2017 Jun;14(6):587–9.

102. Wickham H. ggplot2: Elegant Graphics for Data Analysis. 2nd ed. 2016. Cham: Springer International Publishing : Imprint: Springer; 2016. 1 p. (Use R!).

103. Xu S, Chen M, Feng T, Zhan L, Zhou L, Yu G. Use ggbreak to Effectively Utilize Plotting Space to Deal With Large Datasets and Outliers. Front Genet. 2021 Nov 2;12:774846.

